# Time-resolved and multifactorial profiling in single cells resolves the order of heterochromatin formation events during X-chromosome inactivation

**DOI:** 10.1101/2023.12.15.571749

**Authors:** Samy Kefalopoulou, Pim M. J. Rullens, Kim L. de Luca, Sandra S. de Vries, Tessy Korthout, Alexander van Oudenaarden, Peter Zeller, Jop Kind

## Abstract

The regulation of gene expression is governed at multiple levels of chromatin organization. However, how coordination is achieved remains relatively unexplored. Here we present Dam&ChIC, a method that enables time-resolved and multifactorial chromatin profiling at high resolution in single cells. Analysis of genome-lamina interactions in haploid cells reveals highly dynamic spatial repositioning of small domains during interphase and partial inheritance over mitosis. Dam&ChIC applied to study random X-inactivation uncovers that spreading of H3K27me3 on the inactive X chromosome (Xi) overlaps with remarkable genome-lamina detachment. We find that genome-lamina detachment precedes H3K27me3 accumulation on the Xi and occurs upon mitotic exit. Domains that retain genome-lamina interactions are marked by high pre-existing H3K9me3 levels. These findings imply an important role for genome-lamina interactions in regulating H3K27me3 accumulation on the Xi. We anticipate that Dam&ChIC will be instrumental in unraveling the interconnectivity and order of chromatin events underlying cell-state changes in single cells.

## INTRODUCTION

Gene regulation and the establishment of cellular specificity involves the coordinated activity of processes that take place on different levels of chromatin organization. Such processes include, among others, post-translational modification of histone tails (hPTMs) (Millán-Zambrano et al., 2022), the three-dimensional organization of chromatin (van Steensel & Furlong, 2019) and interactions of chromatin with nuclear scaffolds like the nucleolus and the nuclear lamina (NL) (van Steensel & Belmont, 2017).

Recent advances in molecular technologies have been instrumental for profiling various features of chromatin organization with single-cell precision (Bartosovic et al., 2021; Buenrostro et al., 2015; Kaya-Okur et al., 2019; Kind et al., 2015; Ku et al., 2019; Nagano et al., 2013; Wu et al., 2021; Zeller et al., 2023). Based on these technologies, we learned that the principles of chromatin organization in single cells adhere to findings previously identified by population-based studies, yet they also highlighted that for certain chromatin features, cell-to-cell variability and dynamics are more extensive than previously anticipated.

While methods to measure single-cell chromatin organization are very valuable to reveal heterogeneity, they preclude obtaining insights into the interconnectivity between different chromatin features. To address this, recently several methods were developed to measure multiple chromatin features from the same cell. Experimental approaches such as multi-CUT&Tag (Gopalan et al., 2021), MulTI-Tag (Meers et al., 2022), nano-CT (Bartosovic & Castelo-Branco, 2022), NTT-seq (Stuart et al., 2022) and scMAbID (Lochs et al., 2023) all recover multifactorial chromatin information by using mixtures of target-specific antibodies and *in situ* tagging of the chromatin. The information provided by these methods hold the promise to study cooperativity and the order of gene-regulatory events in the same cell. However, at the same time, the sparsity of the information that most of these technologies provide, makes it challenging to analyze the interplay between chromatin features in single cells. As a result, our understanding of how the different levels of chromatin organization interact and control gene expression in an orchestrated manner remains limited.

Furthermore, the variability and dynamics in chromatin states observed between single cells emphasize the need for tools that can record transitions in chromatin states over time in the same cell. Such technologies would provide unique insights into the role of past chromatin states in determining current cellular outcomes. This time resolution is not feasible with *in situ* tagging approaches because these only capture snapshots of chromatin states, right at the time of harvest. Instead, a time axis could be introduced with a memory system that records information on chromatin-state transitions in living cells, such that they can be retrieved at a later time. Molecular recording is inherent to a few technologies that are based on expression of exogenous bacterial methyltransferases in mammalian cells. When fused to endogenous DNA-interacting proteins, these enzymes can label the genome in live cells, by depositing methylation marks in the proximity of the genomic sites of interaction. These labels represent molecular footprints of past protein DNA-interactions that can be retrieved upon sequencing. This principle was recently illustrated by a technology that is based on the expression of the bacterial cytosine methyltransferase DCM which enabled recordings of gene and enhancer activity during cellular differentiation (Boers et al., 2023). However, since this method was not developed at the single-cell level, it is limited to profiling chromatin-state transitions in populations of cells. Therefore, the necessity to delineate chromatin dynamics that occur within the same cell over time still persists.

A high-resolution single-cell technology that records epigenomic information over a controlled period of time in live cells is (sc)DamID (Guelen et al., 2008; Kind et al., 2015; Rang et al., 2022; Rooijers et al., 2019; van Steensel & Henikoff, 2000; Vogel et al., 2007). DamID involves the fusion of a DNA-binding protein of interest (POI), or a hPTM reader domain, with the E. coli methyltransferase Dam. Conditional expression of the fusion protein in live cells enables the deposition of adenine-6-methylation (m6A) in the proximity of the POI or hPTM. Consequently, protein-DNA contacts or hPTM occupancy are directly recorded on the DNA and remain stable until the DNA is replicated. DamID thereby introduces a time-axis that has the promise to directly measure past chromatin states from the same cell, within a defined window of time. However, information by DamID alone is not sufficient to disentangle past and present chromatin states within the same cell.

In order to develop a multifactorial method that is able to yield time-resolved chromatin states in single cells, here we integrated scDamID (Kind et al., 2015; Rooijers et al., 2019) with the recently developed antibody-based sortChIC method (Zeller et al., 2023). SortChIC recovers high-resolution single-cell chromatin profiles through *in situ* tethering of the non-specific micrococcal nuclease (MNase) fused to protein A (pA-MNase), to specifically digest and amplify antibody-bound chromatin (Schmid et al., 2004; Skene & Henikoff, 2017). Like other *in situ* tagging approaches, sortChIC captures snapshot information on chromatin states at the time of cell harvest. Therefore, our new single-cell method, called Dam&ChIC, combines recording of chromatin states in live cells with antibody-directed chromatin digestion, with a twofold objective: first, to recover multifactorial chromatin information with high resolution to dissect the interplay between chromatin states in single cells; second, to introduce a time-axis in single-cell data to reveal chromatin transitions in the same individual cell.

Here, we demonstrate both properties of Dam&ChIC through comprehensive benchmarking and implementation in two distinct biological systems. First, we provide the proof-of-concept that Dam&ChIC can record chromatin transitions through measuring single-cell dynamics of lamina-associated domains (LADs) during interphase, and the inheritance of LADs through mitosis. We demonstrate that Dam&ChIC can be utilized to obtain structural principles of single-cell LAD-domain reorganizations over time in the same cell and identify a domain-size dependency of LAD inheritance over mitosis. Secondly, we employ Dam&ChIC in X-chromosome inactivation (XCI), an essential part of dosage compensation that depends on heterochromatin formation. We uncover an unanticipated loss of genome-lamina interactions across the Xi that occurs directly prior to accumulation of the heterochromatic mark H3K27me3, an early hallmark event of XCI. Finally, using the time-axis in Dam&ChIC data, we show that the loss of interactions between the Xi and the nuclear lamina occur upon the exit from mitosis. Collectively, we show that Dam&ChIC provides a new experimental and analytical framework that enables studying the ordering of chromatin regulatory events associated with cell-state transitions in single cells.

## RESULTS

### Method design

We integrated scDamID and sortChIC by introducing several adaptations to both protocols, to render them compatible. Dam&ChIC requires creating a cell line that conditionally expresses the Dam enzyme tethered to a protein of interest (POI). First, Dam expression is induced during a desired time window in living cells to deposit m6A on genomic GATC motifs that lie in the proximity of the POI (Fig. 1a, step 1). Cells are subsequently harvested, permeabilized and stained with an antibody of interest (Fig. 1a, step 2). Afterwards, single nuclei are sorted in plates via fluorescence-activated cell sorting (FACS) (Fig. 1a, step 3) and processed with robotic liquid handling using an adapted version of the sortChIC protocol (see Methods). This includes activation of pA-MNase (Fig. 1a, steps 4-5), blunt-ending of the pA-MNase-produced fragments (Fig. 1a, step 6), DpnI digestion to specifically enrich for genomic fragments containing m6A-marked GATC motifs (Fig. 1a, step 7), and ligation with blunt-end forked adapters (Fig. 1a, step 8). The adapter design includes a T7 promoter for *in vitro* transcription (IVT), unique molecular identifiers (UMI), the Illumina P5 sequence and cell-specific barcodes (Rooijers et al., 2019), allowing linear amplification of the produced fragments and illumina library preparation (Fig. 1a, step 9, Fig. S1a). Upon high-throughput sequencing, the unique sequence context of scDamID and sortChIC reads is leveraged to separate the pool of fragments *in silico* (Fig. 1a, step 10, see Methods). Specifically, scDamID reads almost exclusively align to genomic GATC motifs, contrary to sortChIC-derived reads that do not have any motif specificity (Fig. S1b). Due to the intrinsic preference of pA-MNase for A/T-rich genomic regions (Dingwall et al., 1981), more than 95% of sortChIC reads start with either an A or T nucleotide (Fig. S1c-d), which we use to achieve more confident read separation (see Methods). Importantly, the number of reads recovered for both modalities in Dam&ChIC is comparable to control datasets, in which either method is performed individually (control datasets are henceforth referred to as scDamID-only and sortChIC-only) (Fig. S1b, Fig. S1e). In the scDamID-only and sortChIC-only libraries, the vast majority of reads are correctly assigned to the respective readout, separating target from off-target signal by at least two orders of magnitude (Fig. S1e). Therefore, the Dam&ChIC protocol yields sequencing libraries containing both scDamID- and sortChIC-derived fragments, which can be separated *in silico* based on sequence context.

**Figure 1.**
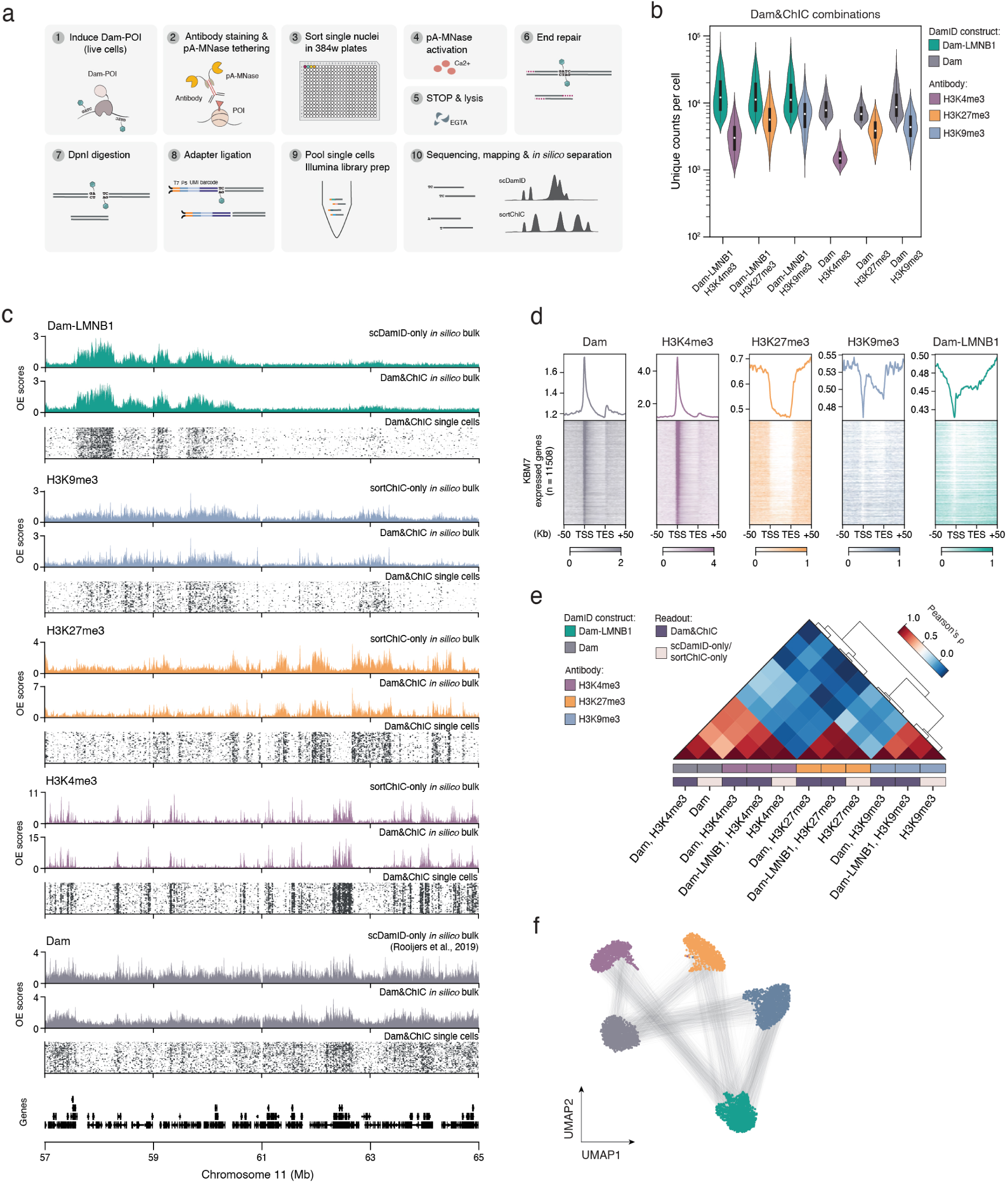
Dam&ChIC enables high resolution joint chromatin profiling in single cells. (**a**) Graphical overview of the single-cell Dam&ChIC method. (**b**) Violin plots depicting the unique number of reads obtained per cell by Dam&ChIC for different combinations of Dam constructs and antibodies. (**c**) Single-cell heatmaps (n = 463-1740 cells) of each chromatin feature profiled with Dam&ChIC and their in silico bulk profiles, shown as observed over expected (OE) values. Additionally, corresponding in silico bulk profiles of sortChIC-only datasets in KBM7 cells are shown for comparison. For the untethered Dam, the in silico bulk of a corresponding publicly available scDamID-only dataset in KBM7 cells (Rooijers et al., 2019) is used for comparison. (**d**) Heatmaps showing chromatin features profiled with Dam&ChIC aligned on genes (n = 11508) expressed in KBM7 cells. Genes (rows) are ordered on their relative expression levels, determined by publicly available RNA-seq data (Rooijers et al., 2019). Profiles above show the scaled averages of the heatmaps for each mapped chromatin feature. (**e**) Hierarchical clustering depicting genome-wide Pearson correlations relating all pseudobulk Dam&ChIC hPTM data and corresponding sortChIC-only datasets, or a publicly available scDamID-only dataset (Rooijers et al., 2019). Data was binned in 1Kb bins. (**f**) UMAP representation of Dam&ChIC data binned in 100Kb genomic bins. Each cell (dot) is represented twice, once for each measurement. Black lines connect the same cell.

### Dam&ChIC jointly maps diverse chromatin types with high resolution

To benchmark the quality of the integrated protocol, as well as to test its versatility in profiling different chromatin features, we generated single-cell data of diverse combinations of heterochromatic and euchromatic chromatin types. We used two previously established human KBM7 cell lines; one that conditionally expresses Dam tethered to the core nuclear lamina protein LMNB1 and one that conditionally expresses the untethered Dam enzyme. Dam-LMNB1 has been previously used to characterize LADs in single cells (Kind et al., 2015), while the untethered Dam enzyme has been reported to accurately detect chromatin accessibility in single cells (Rooijers et al., 2019). Importantly, KBM7 cells have a near complete haploid genome, ensuring that both Dam&ChIC measurements originate from the same chromosome copy. For these experiments, we induced expression of Dam-LMNB1 or untethered Dam in live cells for 15 hours, to enable m6A deposition. Thereafter, we stained nuclei with antibodies specific to the hPTMs H3K4me3, H3K27me3 or H3K9me3. For benchmarking, we additionally performed sortChIC-only experiments for the same set of hPTMs and used matching scDamID-only data, both derived from KBM7 cells.

For Dam-LMMB1 and the untethered Dam, Dam&ChIC recovers a median of ∼12,300 and ∼8,500 UMI-flattened (unique) reads per cell respectively, in all combinations with hPTMs (Fig. 1b). For the euchromatic H3K4me3 it recovers a median of ∼2,600 unique reads per cell and for the heterochromatic H3K27me3 and H3K9me3 modifications a median of ∼5,000 and ∼5,500 unique reads per cell respectively (Fig. 1b). Dam&ChIC thereby attains higher sensitivity compared to other recently published multifactorial chromatin profiling methods, particularly considering the haploid genome of KBM7 cells (Fig. S1f). In order to normalize the two distinct fragment types present in the Dam&ChIC libraries, we computed observed over expected (OE) scores (Kind et al., 2015). For the scDamID readout, OE scores are calculated over the in silico genomic distribution of GATC motifs (Fig. S1g, see Methods). For the sortChIC readout, the distribution of maximum expected reads was generated with a bulk sortChIC-only experiment against H3 (Fig. S1g, see Methods).

We directly compare Dam&ChIC to sortChIC-only and scDamID-only data, at the resolution of individual genes and lamina-associated domains (LADs). This confirms highly specific single-cell genomic enrichment at the expected regions (Fig. 1c,d, Fig. S1h), further corroborated by a comparison against matching ENCODE bulk data of the closely related K562 cell line (Fig. S1i,j). Moreover, a genome-wide comparison shows strong correlation between Dam&ChIC and the corresponding sortChIC-only and scDamID-only datasets (Fig. 1e). Finally, dimensionality reduction of the entire Dam&ChIC dataset reveals that the different single-cell readouts consistently separate based on chromatin type (Fig. 1f). Altogether, these results demonstrate that Dam&ChIC is a versatile multifactorial method for profiling diverse euchromatic and heterochromatic factors in single cells with high sensitivity and specificity.

### Dam&ChIC disentangles past and present genome-lamina interactions in the same cell

Using a fluorescent tracker to follow Dam-labeled DNA by microscopy (m6A-Tracer) we previously reported that, during interphase, LADs display constrained localization dynamics within a 1 um zone underneath the nuclear lamina (Kind et al., 2013). These findings indicate that over a 15-hour time window LADs are very dynamic, and that these dynamics can be efficiently detected with DamID (Kind et al., 2013). In contrast, the sortChIC readout can only detect LADs that are in contact with the nuclear lamina at the time of cell harvest. To explore the possibility to use the Dam&ChIC method to capture temporal dynamics of LADs during interphase, we used the Dam-LMNB1 KBM7 cell line to induce expression of the Dam-LMNB1 fusion protein for a period of 15 hours, during which genome-lamina interactions are directly recorded on the DNA. After this period, the cells were stained with an antibody against LMNB1 to detect current-state genome-lamina interactions with the sortChIC readout. Provided that genome-lamina interactions undergo a degree of dynamics in KBM7 cells, we expect that past interactions will be detected exclusively by scDamID (Fig. 2a; past interactions), whereas very recently established interactions are only measured with sortChIC (Fig. 2a; de novo interactions).

**Figure 2.**
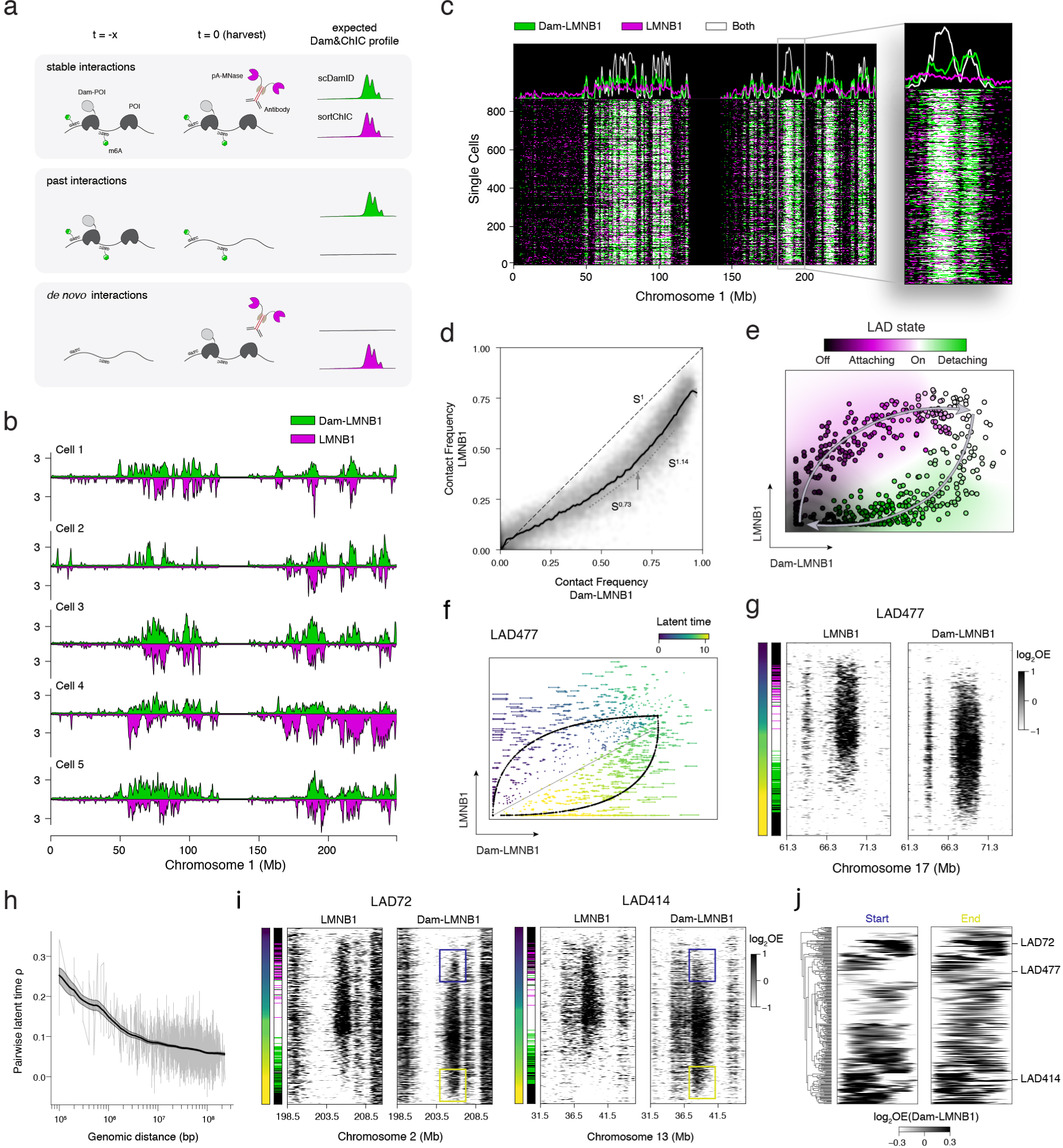
Dam&ChIC disentangles past and present genome-lamina interactions in the same cell. (**a**) Schematic overview of the chromatin dynamics that Dam&ChIC can resolve between the time of Dam-POI induction (t = -x) and the time of cell harvest for antibody staining (t = 0). Three scenarios are depicted with their corresponding expected Dam&ChIC profiles. (**b**) “Mirror plots” showing quantitative LAD signal in 100Kb genomic bins on chromosome 1 measured by scDamID (top, green) and sortChIC (bottom, magenta) within the same cell for 5 example cells. (**c**) Combined single-cell heatmap (n = 865) with binarized Dam&ChIC LAD signal (see Methods) in 100Kb genomic bins on chromosome 1. Bins are detected by either scDamID (green), sortChIC (magenta), or by both measurements (white). (**d**) Scatter plot comparing the contact frequency (CF) (see Methods) of the scDamID (x-axis) and sortChIC (y-axis) measurement of Dam&ChIC. Black uninterrupted line indicates a radius neighbor regression model that was fit on the data, with its 95% confidence interval. Diagonal black dashed line indicates complete agreement between both measurements. Slope is quantified at different positions. (**e**) Schematic of scVelo model (Bergen et al., 2020) implementation that describes dynamics of an individual LAD based on the ratio of Dam&ChIC LMNB1 data (scDamID: x and sortChIC: y axes). Each dot is a cell for which the state of a given LAD is colored according to the likelihood of being off (black), attaching to the lamina (magenta), on (white) or detaching from the lamina (green). (**f**) Phase portrait of LAD477 inferred by scVelo, with arrows indicating velocities of single cells colored according to latent time. Curved lines indicate the velocity model and straight line the steady state. (**g**) Single-cell heatmaps for LAD477, with the scDamID (right) and sortChIC (left) measurements split and single cells sorted according to LAD477-based latent time (leftmost bar). The middle bar at the left indicates the LAD state (see Methods). LAD signal is shown as log-trans-formed observed over expected (OE) in 100Kb genomic bins. (**h**) Pairwise Pearson’s correlation coefficient of the latent time assignment between any pair of LADs (y-axis), plotted against genomic distance (x-axis). Black line indicates a radius neighbor regression model that was fit on the data, with its 95% confidence interval. (**i**) Single-cell heatmaps of two other example LADs (LAD72, left and LAD414, right) and their dynamics, visualized like in (g). Blue and yellow boxes indicate the origin and release of each domain respectively, happening from and towards the same genomic location. (**j**) Hierarchical clustering of latent time averaged start (left) and end (right) of LADs at which they emerge or disappear. Latent time averages are from 50 cells up and downstream of start or end, as depicted in blue and yellow boxes, respectively, in 2i. LAD signal is shown as log-transformed observed over expected (OE) in 100Kb genomic bins.

We obtained high quality single-cell LAD profiles for both Dam&ChIC measurements (Fig. S2a-b). We started by examining – on a population level – which LADs are identified by both measurements, or by scDamID or sortChIC alone. For this purpose, we calculated the contact frequency (CF) metric that quantifies the fraction of cells in which a genomic region is in contact with the nuclear lamina (Fig. S2b; see Methods) (Kind et al., 2015). We find a strong overlap between LAD profiles identified by scDamID and sortChIC, with only a small minority of LADs detected exclusively by one of both measurements (Fig. S2c). Moreover, the CF is comparable between both measurements, albeit somewhat higher for several regions in the scDamID measurement (Fig. S2c).

We next inspected the combined genome-lamina contact profiles in single cells. Interestingly, visual inspection of Dam&ChIC data in single cells indicates that although LADs are generally detected by both methods, we also regularly observe LADs that are identified exclusively by scDamID and to a lesser extent by sortChIC alone (Fig. 2b). This was confirmed by systematic analysis of the fraction of 100 kb genomic bins that are identified only by scDamID, sortChIC, or by both (Fig. 2c, Fig. S2d-f; see Methods). Notably, the genomic bins that display the largest discrepancy between scDamID and sortChIC signal are the genomic regions that interact with the nuclear lamina with intermediate CFs which indicates that these regions display the highest dynamics during interphase (Fig. 2d). In contrast, regions with higher CFs are more comparably detected by both Dam&ChIC readouts, suggesting more stable genome-lamina interactions for these regions, that typically reside in larger domains with high A/T content (Fig. 2d) (Kind et al., 2015). This finding discerns variable degrees of dynamics between different regions that interact with the nuclear lamina and corroborates our previous observations that larger LADs with high CFs more stably associated with the nuclear lamina through multivalent interactions over hundreds of kilobases (Kind et al., 2015).

Given that the current Dam&ChIC dataset captures genome-lamina dynamics, we reasoned that the ratio between scDamID and sortChIC signal in a given cell should contain information on the direction of LAD changes. High sortChIC and low scDamID signal indicates that a LAD is being established. Conversely, low sortChIC and high scDamID signal implies that a LAD is being released from the nuclear lamina. Based on this reasoning, we next sought to derive structural principles in LAD-domain establishment and release.

RNA velocity (La Manno et al., 2018) and the ensuing chromatin velocity (Tedesco et al., 2022) are based on estimating the ratio between two interdependent cellular modalities to extract directional change. We apply this principle to Dam&ChIC and infer the chromatin velocity of genome-lamina interactions, based on the ratio between scDamID and sortChIC (Fig. 2e). This results in a LAD-specific latent time (Bergen et al., 2020), that orders each LAD according to its state of establishment and release over time (Fig. 2f,g, Fig. S2g). In agreement with previous findings that LADs show coordinated nuclear lamina association along the linear chromosome (Kind et al., 2015), we find that nearby LADs progress coordinately, with a decay in their coordination according to linear intra-chromosomal distance (Fig. 2g,h). We further observe remarkable polarity in the establishment of some LADs; one border of the domain is visible first and defines from where the domain will expand to the opposing border (Fig. 2i). Interestingly, release of the domain from the nuclear lamina mirrors the patterns of LAD establishment (Fig. 2i,j). Regardless of the position where the domain appears first (i.e., left, central or right), LADs generally start narrow and broaden from this region, to be released from the nuclear lamina at the position of initial appearance (Fig. 2g,i,j).

Collectively, we demonstrate that Dam&ChIC disentangles past and present chromatin states in the same cell. Our analyses reinforce and extend upon previous findings of genome-lamina interaction dynamics (Kind et al., 2013) and reveal that the degree of LAD dynamics in interphase is linked to the CF of a given region in a cell population. By interpreting the ratio between the two Dam&ChIC measurements, we demonstrate that Dam&ChIC can be used to derive structural principles of chromatin domain dynamics.

### Spatial positioning of the genome is partially inherited upon mitosis

Encouraged by the observations that Dam&ChIC disentangles past from present chromatin states, we set out to implement it to study the inheritance of spatial genome organization over a cell division. Previous work described differences in genome-lamina interactions between mother and daughter cells, as a result of stochastic re-shuffling of LADs right after mitosis (Kind et al., 2013). These observations were made by microscopy using the m6A-Tracer, which enabled imaging of LAD inheritance, but left sequence identity and the underlying patterns of genome-lamina reorganization elusive. In order to address this in a genome-wide manner, we sought to leverage Dam&ChIC to measure genome-lamina interactions before and after mitosis in the same cell.

To this end, we synchronized KBM7 cells in G1/S using a double thymidine block and expressed Dam-LMNB1 concurrently with the last thymidine incubation (Fig. S3a and Methods). We harvested cells in G2 phase and early G1 phase (Fig. S3a and Methods) and utilized a multiplexing sorting strategy (Yeung et al., 2022) to enable parallel processing of different conditions with Dam&ChIC (Fig. S3a-b and Methods). Of note, while the Dam-deposited m6A mark is typically not propagated upon DNA replication in the absence of the Dam-POI, our experimental setup ensures that Dam-LMNB1 is still present during S and G2 in order to label and record G2 LADs (Fig. S3a and Methods).

In G2 cells, Dam&ChIC is expected to yield extensive similarity between scDamID and sortChIC, because of the narrow window of time between m6A deposition and antibody staining. Importantly, G2-cell profiles can serve as a baseline control to contrast any changes in genome-lamina interactions that occur upon cell division in early G1. In early G1 cells, LADs that are faithfully inherited through mitosis will be detected both by scDamID and sortChIC (Fig. 3a, cell 1, domains a, b, c). In contrast, LADs that reposition away from the nuclear lamina after mitosis are exclusively recovered by scDamID (Fig. 3a, cell 2, domains a, c), whereas de novo established LADs will be exclusively recovered by sortChIC (Fig. 3a, cell 2, domain d).

**Figure 3.**
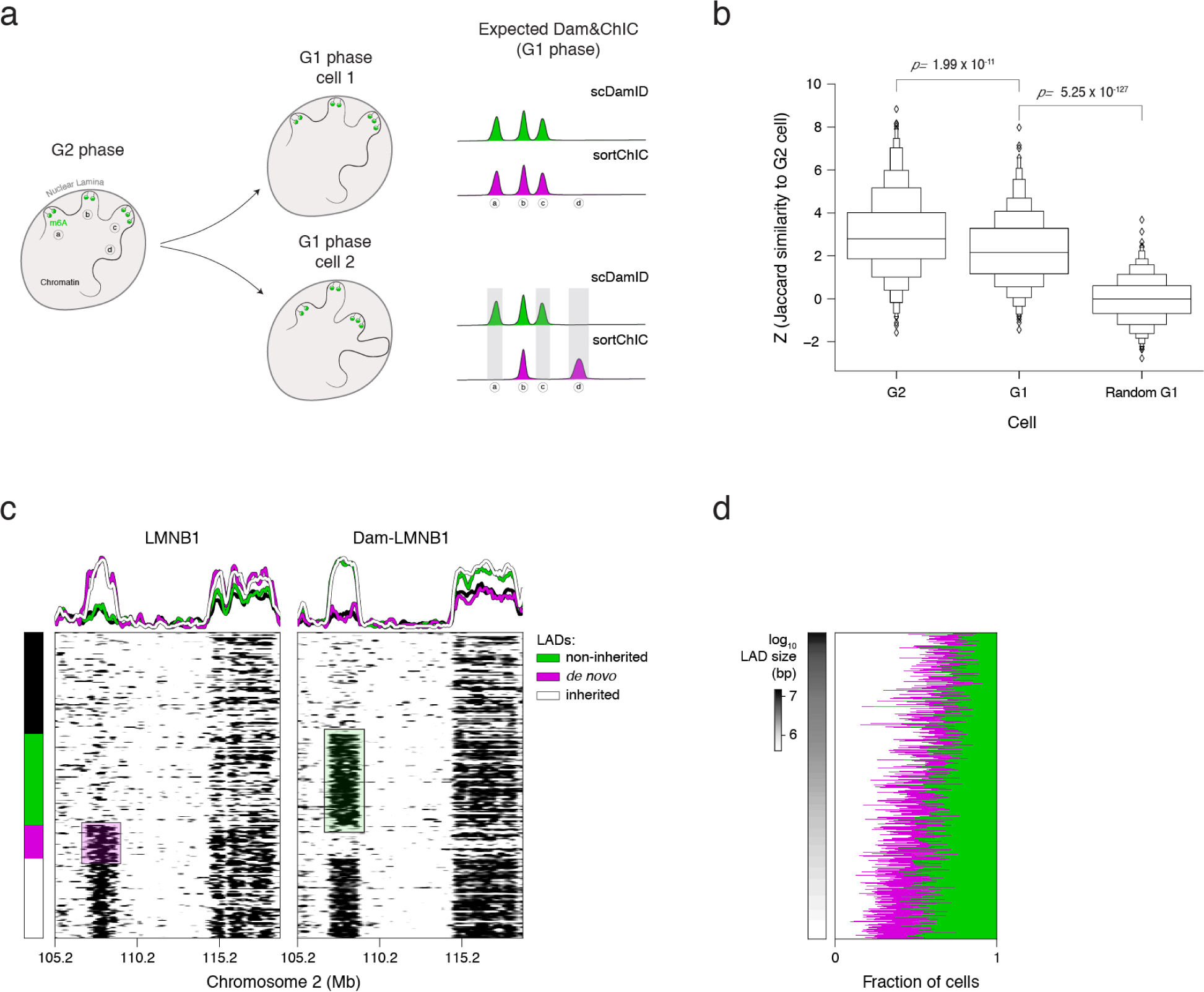
Dam&ChIC identifies principles of genome-lamina inheritance upon mitosis. (**a**) Schematic of experiment and hypothetical outcome of LAD inheritance upon cell division measured by Dam&ChIC. LADs that are inherited to cell 2 (b) are expected to be captured in G1-phase by both scDamID and sortChIC. LADs non-inherited to cell 2 (a,c) are expected to be recovered only by scDamID, while de novo LADs of cell 2 (d) to be recovered only by sortChIC in G1 phase cells. (**b**) Pairwise z-score normalized Jaccard similarity between scDamID of G2 cell and sortChIC of the same G2 cell (left), descending G1 cell (middle) or random G1 cell (right). (**c**) Single-cell heatmaps of scDamID and sortChIC measurements of Dam&ChIC, showing two LADs on Chromosome 2 that are either weakly inherited (left) or faithfully inherited (right). Cells are grouped according to the recovery of the left LAD by scDamID and/or sortChIC as non-inherited, de novo or inherited. (**d**) Quantification of the fraction of cells for which a given LAD is found to be non-inherited, de novo or inherited, ordered according to LAD size (n = 516).

We first confirm that cells in G2 display typical LAD profiles (Fig. S3c), verifying that our synchronization and induction approach does not introduce adverse effects on genome-lamina interactions. We then compute the z-score normalized Jaccard index, to measure the pairwise similarity between scDamID and sortChIC measurements (see Methods) for three pairs: (i) between scDamID and sortChIC of the same cell in G2, (ii) between scDamID and sortChIC of the same cell in early G1 (that is, scDamID patterns of G2 before cell division, and sortChIC of early G1 after cell division) or (iii) between scDamID and sortChIC of random cells in G1. We observed a significant decrease in similarity between scDamID and sortChIC LAD profiles in G1 compared to G2 cells, yet, the similarity in G1 cells is considerably higher than expected by random chance (Fig. 3b). This suggests that LADs are partially inherited over mitosis, corroborating previous microscopy findings (Kind et al., 2013). Interestingly, we can accurately detect three scenarios: inherited LADs in G1 (Fig. 3c; white), de novo established LADs in G1 (Fig. 3c; purple) and non-inherited LADs, unique to G2 cells (Fig. 3c; green). The occurrence of these scenarios differs extensively, with some domains showing considerable heterogeneity in inheritance across cells (Fig. 3c; left domain), while others are stably preserved across generations (Fig. 3c; right domain).

We next wondered if some LADs have a higher propensity to be inherited through mitosis than others. Interestingly, we find that larger LADs are more likely to be inherited compared to small LADs (Fig. 3d). This is further corroborated by an increased similarity in the G1 population for large LAD sizes, compared to small LAD sizes (Fig. S3d). This data indicates that larger LADs tend to be stable during G2 and faithfully inherited through mitosis, in contrast to smaller LADs that are more dynamic during G2 and less frequently inherited.

In summary, this dataset shows the unique ability of Dam&ChIC to capture chromatin transitions over a cell division. Additionally, it provides an experimental and analytical framework to obtain insights into the principles of epigenetic inheritance of spatial genome organization.

### Dam&ChIC reveals widespread remodeling of genome-lamina interactions during X-chromosome inactivation

Above we established that Dam&ChIC, as a multifactorial method, can profile diverse chromatin features at high single-cell resolution. This ability makes Dam&ChIC an ideal approach to study the interplay between chromatin features in dynamic cellular settings that undergo multiple chromatin transitions over time. An essential process during mammalian development that entails dynamics of multiple chromatin features, is X-chromosome inactivation (XCI). XCI is specific to female cells to ensure dosage compensation of X-linked genes between males (XY) and females (XX) (Lyon, 1961; Lyon, 1962), through the formation of transcriptionally repressive heterochromatin on the inactive X chromosome (Xi) (Heard et al., 2004).

Heterochromatin formation on the Xi is regulated primarily by the long non-coding RNA Xist (Brown et al., 1992; Panning & Dausman, 1997; Penny et al., 1996) and subsequent deposition and spreading of H3K27me3 (Plath et al., 2003; Silva et al., 2003; Żylicz et al., 2019) and other repressive hPTMs (reviewed by Zylicz & Heard, 2020). While remodeling of the hPTM landscape on the Xi has been under continuous investigation, the nuclear positioning of the Xi remains unclear. Previous studies suggest that the Xi associates with different nuclear compartments, such as the nucleolus or the nuclear lamina (NL) (Belmont et al., 1986; Bourgeois et al., 1985; Chen et al., 2016; Rego et al., 2008; Teller et al., 2011; Zhang et al., 2007). Whether this interaction is dynamic and how it is linked to other hPTMs during XCI has remained ambiguous. Moreover, the random nature of XCI has forced population-based genomic studies to work with genetically engineered hybrid systems that predetermine which of the two X alleles will be inactivated (Augui et al., 2011), making it challenging to decouple observations on XCI from potential biases introduced by the genetic background.

With Dam&ChIC, we aimed to map the interactions of the Xi with the nuclear lamina and dissect the interplay with hPTMs H3K27me3 and H3K9me3. To this end, we induced random XCI *in vitro*, by differentiating our two previously derived female hybrid (CAST/Eij x 129/Sv) embryonic stem cell (ESC) lines expressing Dam-LMNB1 and Dam-scFv-H3K27me3 (Guerreiro et al., 2023; Rang et al., 2022) (Fig. 4a). We used an established *in vitro* differentiation protocol that involves Vitamin C treatment on a monolayer of cells and attains asynchronous but robust inactivation within a few days (Loda et al., 2017; Robert-Finestra et al., 2021). We performed Dam&ChIC at multiple timepoints between day 0 and 6 of differentiation, using the same multiplexing sorting strategy described earlier (see Methods). We compiled dual measurements of Dam-LMNB1/H3K27me3 (n = 1656 cells), Dam-LMNB1/H3K9me3 (n = 573 cells), Dam-scFv-H3K27me3/LMNB1 (n = 921 cells) and Dam-scFv-H3K27me3/H3K9me3 (n = 497 cells) (Fig. 4a). Upon allelically resolving the Dam&ChIC data based on single nucleotide polymorphisms (see Methods), we consistently detect a high number of unique counts per allele for all measurements (Fig. S4a), providing ample information to interrogate the interplay between the chromatin features of interest.

**Figure 4.**
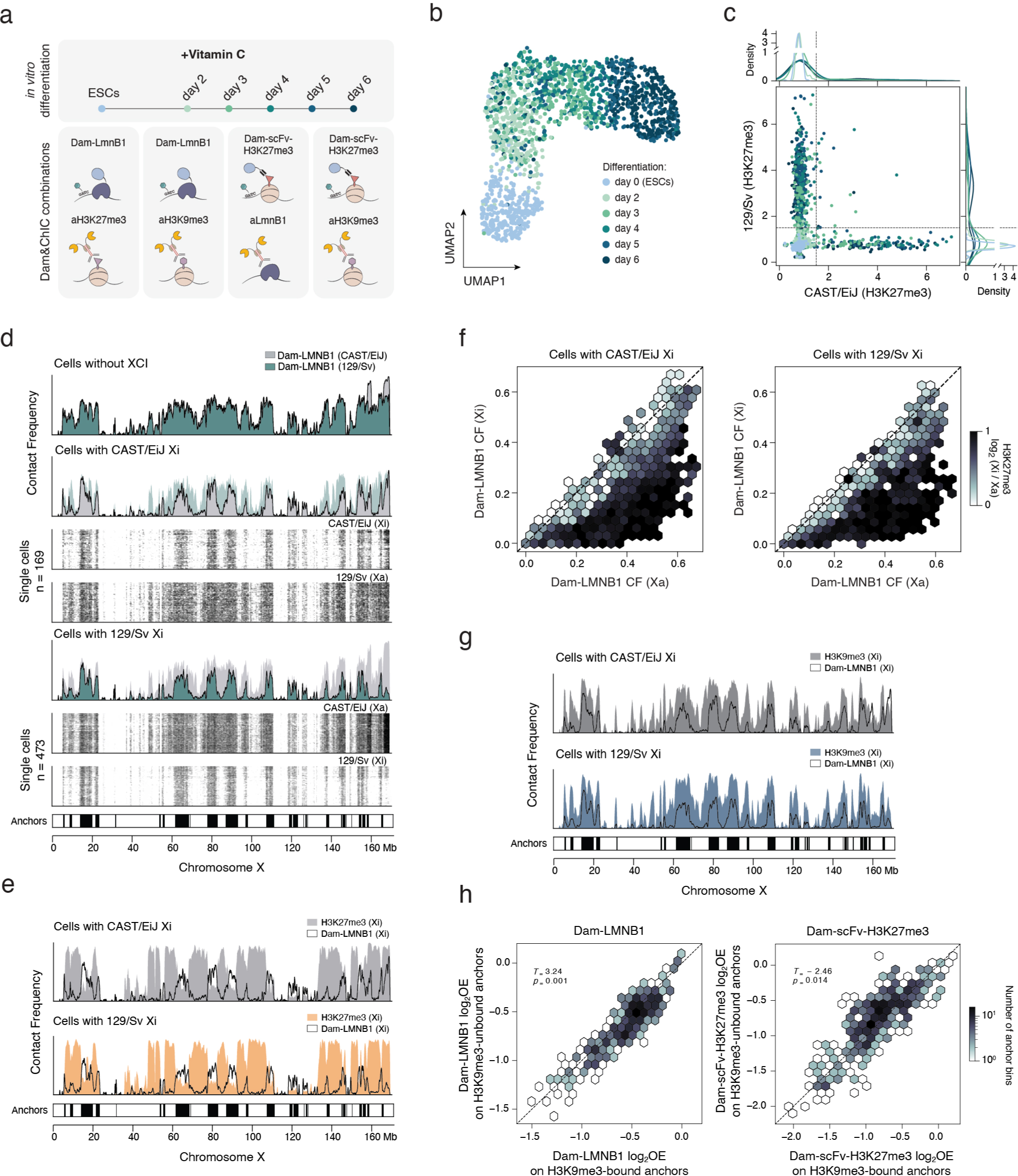
Dam&ChIC reveals widespread genome-lamina detachment during X chromosome inactivation. (**a**) Schematic of the experimental setup with the Vitamin C differentiation trajectory and the Dam&ChIC combinations. (**b**) UMAP based on H3K27me3 levels on autosomal genes of Dam-LMNB1/H3K27me3-profiled cells, colored by differentiation day. (**c**) Scatterplot of allelic X-chromosomal H3K27me3 levels per single cell in the Dam-LMNB1/H3K27me3 dataset, colored by differentiation day. The dashed lines indicate the manually set threshold used to classify cells as “no XCI”, “CAST/EiJ inactive”, “129/Sv inactive” or “undetermined”. (**d**) Contact Frequency values of Dam-LMNB1 on either allele for the cell categories defined in (4c), along the entire X chromosome. For the categories that undergo XCI, single-cell heatmaps of the active (Xa) and inactive (Xi) X chromosome are shown below, with the respective cell numbers. Bottom bar shows calls of the “anchors”. (**e**) Contact Frequency values of Dam-LMNB1 and corresponding H3K27me3 values on either allele in the inactive state for the cell categories defined in (4c) along the entire X chromosome. Bottom bar shows calls of the “anchors”. (**f**) Hexbin plot of Dam-LMNB1 Contact Frequency (CF) values in cells where the indicated allele is inactivated (left:CAST/EiJ, right:129/Sv). For each LAD the average CF is shown for the active (Xa) versus the inactive (Xi) allele Hexbins are colored based on the log-transformed enrichment of H3K27me3 (Xi/Xa). (**g**) Contact Frequency values of Dam-LMNB1 and H3K9me3 on either allele in the inactivated state, similar to (e). (**h**) Comparison of Dam-LMNB1 (left) and Dam-scFv-H3K27me3 (right) levels between cells in which a specific nuclear lamina “anchors” is H3K9me3 bound (x-axis) or unbound (y-axis). Hexbins are coloured by anchor abundance. Two-sided T-test with resulting p-values are designated.

First, we examined the Dam-LMNB1/ H3K27me3 dataset. Apart from dissecting its interplay with genome-lamina interactions, the H3K27me3 readout can serve two additional purposes: (i) as a means to estimate the differentiation progression, and (ii) as a proxy for the progression of XCI. We used H3K27me3 enrichment on autosomal genes to visualize cells by UMAP, and observe gradual transitioning of cells over differentiation time (Fig. 4b). Additionally, we observe strong H3K27me3 enrichment either on the CAST/Eij or 129/Sv X-chromosome allele over the course of differentiation (Fig. 4c), which we subsequently use to categorize cells as CAST/ Eij-inactive, 129/Sv-inactive or cells that have yet to undergo XCI (Fig. S4b-c). The number of cells that undergo XCI increases with each day of differentiation, with a clear preference for the 129/ Sv allele (Fig. 4c, Fig. S4b).

We next compare the genome-lamina interactions of the X chromosome across the H3K27me3-derived categories (i.e., no XCI, CAST/Eij inactive or 129/Sv inactive). In contrast to the suggested perinuclear localization of the Xi (Chen et al., 2016), we find widespread loss of interactions between the Xi and the nuclear lamina (Fig. 4d, Fig. S4d). This is not the case for its active counterpart (Xa), which retains genome-lamina interactions similar to cells that did not undergo XCI (Fig. 4d, Fig. S4d). Reassuringly, we find the exact same patterns in the inverse experiment where H3K27me3 is measured with scDamID and LADs with sortChIC, confirming that the observed loss in genome-lamina interactions is unlikely a technical artifact (Fig. S4e-g). The loss of interactions between the Xi and the nuclear lamina extends through megabase-sized regions across the entire X chromosome and is interspersed with regions that maintain strong interactions (henceforth referred to as “anchor” regions). Upon cross-examining the loss of genome-lamina interactions and the increase of H3K27me3 on the Xi, we find that regions with lost interactions are densely covered with H3K27me3, as opposed to the anchor regions that are largely devoid of this mark (Fig. 4e). In addition, we quantitatively corroborate the mutual exclusivity between genome-lamina interactions and H3K27me3 on the Xi, showing that the regions that lose lamina interactions upon XCI specifically gain H3K27me3 (Fig. 4f).

Genomic regions with constitutive genome-lamina interactions are generally enriched for the repressive hPTMs H3K9me2/3 (Guelen et al., 2008). We therefore wondered if the resistance of the Xi anchor regions to detachment from the nuclear lamina and H3K27me3 deposition might be related to the presence of H3K9me3. Indeed, Dam&ChIC datasets profiling Dam-LMNB1/ H3K9me3 and Dam-scFv-H3K27me3/H3K9me3 indicate that H3K9me3 is specifically enriched at the anchor regions of the Xi (Fig. 4g), which is consistent with previous observations that H3K27me3 and H3K9me3 occupy distinct non-overlapping domains along the Xi (Vallot et al., 2015). Intriguingly, H3K9me3 domains preexist in cells that did not yet undergo XCI, where they already demarcate the future anchor regions (Fig. S4h). Similar to small LADs on autosomes, individual anchor regions are not present in every single cell. We therefore wondered whether these cell-to-cell differences in anchor usage might be linked to the pre-existing H3K9me3 state. To this end, we quantified the genome-lamina interactions and H3K27me3 deposition at anchors that are either H3K9me3 bound or H3K9me3 unbound across single cells. We find that anchors that are H3K9me3 bound in individual cells are more likely to interact with the nuclear lamina and less likely to be enriched with H3K27me3 (Fig. 4h). These findings suggest that lamina-interacting regions on the X chromosome that are devoid of H3K9me3 enrichment, are predetermined to detach from the nuclear lamina and accumulate H3K27me3 during XCI.

In summary, Dam&ChIC provides allelically-resolved single-cell maps of genome-lamina interactions and heterochromatic hPTMs from the same cell. We find that large X-chromosome regions detach from the nuclear lamina during mouse XCI, and instead become strongly enriched for H3K27me3. Regions that are resistant to detachment and H3K27me3 deposition coincide with pre-existing H3K9me3 domains.

### The Xi detaches from the nuclear lamina prior to H3K27me3 accumulation

The observation that genome-lamina interactions and H3K27me3 accumulation show a mutually exclusive pattern on the Xi prompted us to examine the temporal order between these two seemingly antagonistic events. To identify the onset of nuclear lamina detachment and H3K27me3 accumulation at single-cell resolution, we first ordered cells along the differentiation trajectory by inferring pseudotime based on the UMAP presented in Fig. 4b (Cao et al., 2019; Trapnell et al., 2014) (Fig. 5a). Of note, this UMAP is based solely on H3K27me3 signal on autosomal genes, leaving out the X-linked genes, to avoid skewing the temporal ordering of cells by the H3K27me3 accumulation on the Xi. Interestingly, the initial loss of genome-lamina interactions appears to coincide with the emergence of H3K27me3 around day 3 of differentiation, further implying their interrelatedness (Fig. 5b). We observe a seemingly concurrent onset of both events when visualizing the genomic distribution of genome-lamina interactions and H3K27me3 on the Xi for progressive pseudobulk profiles (Fig. 5c) and single cells (Fig. 5d).

**Figure 5.**
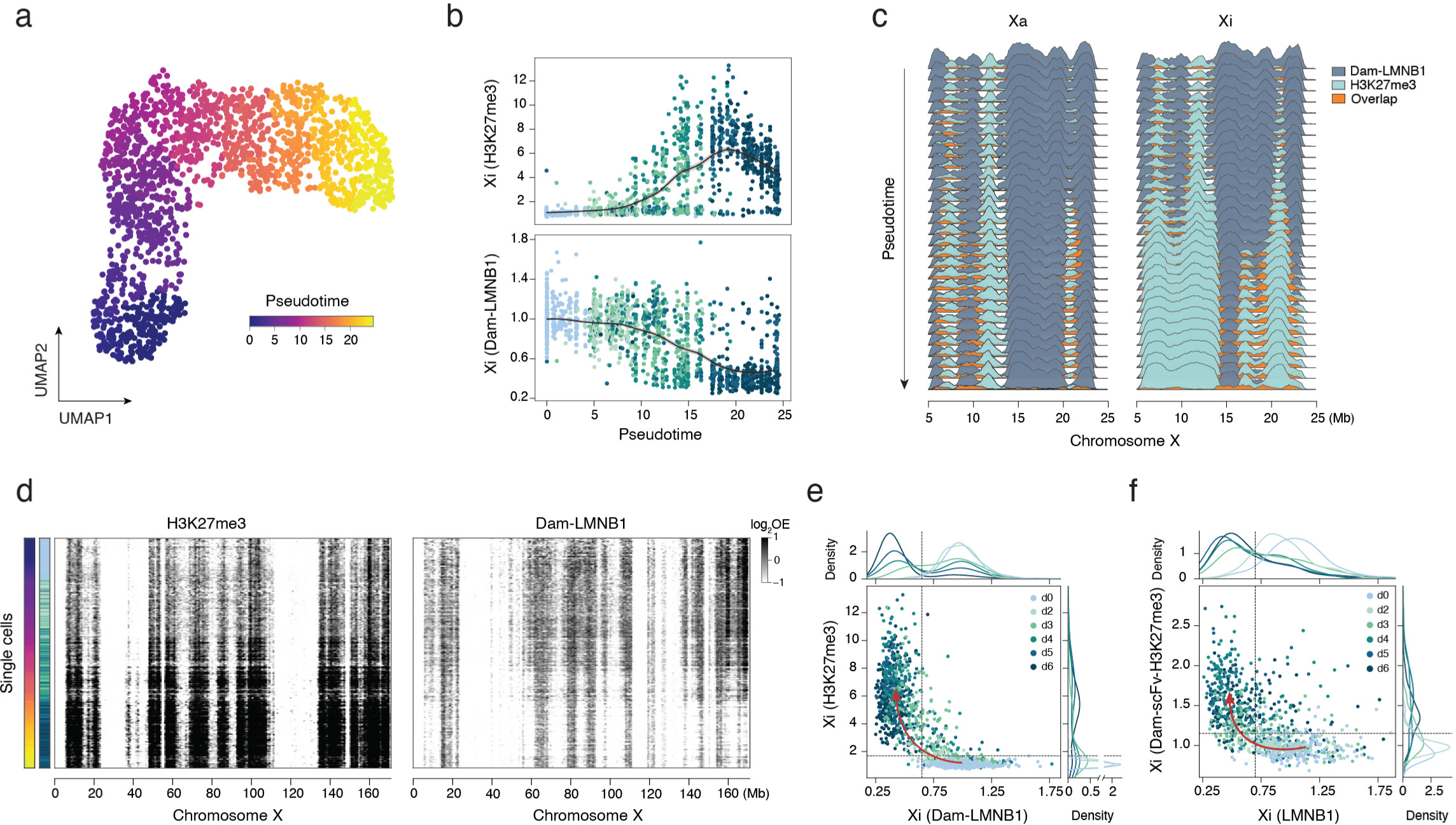
Dam&ChIC resolves order of genome-lamina detachment and H3K27me3 X-chromosome inactivation events. (**a**) Same UMAP as in (4b) based on H3K27me3 levels on autosomal genes, but colored by Monocle3-inferred pseudotime (Cao et al., 2019). (**b**) The inactive X (Xi) average H3K27me3 (top) and Dam-LMNB1 (bottom) levels (y-axis) across pseudo-time (x-axis) are shown. Cells are colored according to differentiation day. (**c**) Contact Frequency of Dam-LMNB1, H3K27me3 and their overlap for groups of 100 cells progressively slid along pseudotime (y-axis) for a dynamic region of the active X chromosome (Xa; left) and its inactive counterpart (Xi; right). (**d**) Single-cell heatmaps ordered along pseudotime (y-axis) of the entire inactive X-chromosome (x-axis) for H3K27me3 (left) and Dam-LMNB1 (right). (**e**) Scatterplot of the average inactive X (Xi) Dam-LMNB1 (x-axis) and H3K27me3 (y-axis) levels. Cells are colored according to differentiation day. Distri-butions above and on the right show average density per day. Arrow indicates directionality of cells. (**f**) Same as (e), but with the inverse readout: Dam-scFc-H3K27me3 (y-axis) / LMNB1 (x-axis).

Therefore, to resolve the precise chronology between nuclear lamina detachment and H3K27me3 enrichment, we plot Dam-LMNB1 against H3K27me3 levels on the Xi for each cell in our dataset, and confirm the inverse relationship between H3K27me3 and genome-lamina interactions at the single cell level (Fig. 5e). Interestingly, early time points show a reduction in Dam-LMNB1 levels before H3K27me3 starts to emerge, resulting in a general left-upward mobility of cells over differentiation in the Dam-LMNB1 to H3K27me3 space (Fig. 5e, arrow). This observation suggests that the Xi first loses its genome-lamina interactions before accumulating H3K27me3. We anticipate that the opposite order of events would result in the temporal appearance of double positive cells and a general upward-right mobility of cells over differentiation. To corroborate our reasoning, we perform simulations where we control the temporal ordering of the two dynamics (Fig. S5a-c), and confirm that the observed left-upward mobility in Fig. 5e corresponds to the simulation where loss of genome-lamina interactions occurs prior to H3K27me3 accumulation. Importantly, we identify the same pattern in the inverse Dam&ChIC experiment profiling Dam-scFv-H3K27me3/ LMNB1, which detects an even earlier loss of genome-lamina interactions (Fig. 5f). These observations collectively support a model in which the loss of genome-lamina interactions is an early event in mouse XCI that precedes, and potentially regulates, H3K27me3 accumulation on the Xi.

### Nuclear lamina detachment of the Xi occurs upon mitotic exit

We earlier demonstrated that principles of chromatin velocity can be used on Dam&ChIC data to resolve the directional dynamics of chromatin domains. We next aimed to extend this principle to XCI, to obtain further insight into the timing and dynamics of Xi’s localization in the nucleus. To this end, we performed an additional Vitamin C differentiation experiment where we mapped genome-lamina interactions with both scDamID and sortChIC readouts (Dam-LMNB1/ LMNB1). By directly estimating the ratio between scDamID and sortChIC, we infer the velocity of genome-lamina interactions of the Xi to identify the moment of detachment and examine its link to the underlying cell state during XCI.

First, based on Xi Dam-LMNB1 signal we visualize cells by UMAP and observe gradual transitioning of cells during differentiation (Fig. 6a). The inferred chromatin velocity accurately recapitulates the direction of differentiation and change in genome-lamina interactions during XCI (Fig. 6a), and reassuringly, fails to deduce any directionality when the ratio between scDamID and sortChIC of randomly coupled cells is used (Fig. S6a). Strikingly, while UMAP does not position undifferentiated ES cells at the start of the trajectory, the inferred chromatin velocity consistently identifies them as root cells without *a priori* designation. This accurate temporal ordering is also reflected in the inferred latent time, that is purely grounded on genome-lamina interaction dynamics, in contrast to similarity-based pseudotime (Fig. 6b, Fig. S6b-c). Moreover, latent time resolves the detachment process during XCI (Fig. 6c, Fig. S6d).

**Figure 6.**
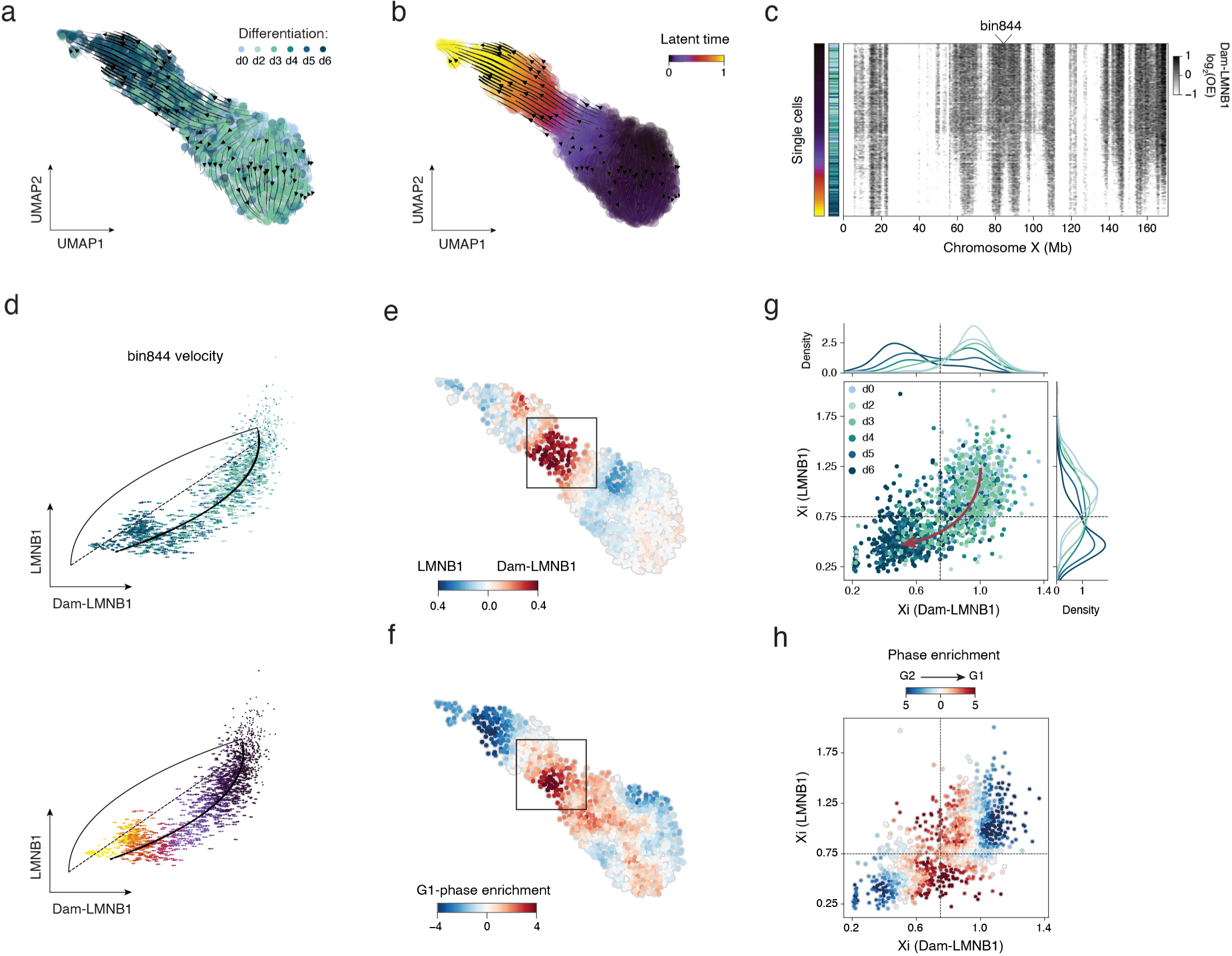
Inactive X genome-lamina velocity identifies cell division-dependent detachment. (**a**) UMAP based on Dam-LMNB1 levels of the inactive X 100Kb bins colored according to differentiation day. Streamlines on top of UMAP represent chromatin velocities inferred by the ratio of Dam-LMNB1 and LMNB1 100Kb inactive X bins, using scVelo (Bergen et al., 2020). (**b**) Same UMAP as (a), but with cells colored according to scVelo inferred latent time, that is purely based on Dam-LMNB1/LMNB1 dynamics, not on the underlying UMAP. (**c**) Dam-LMNB1 single-cell heatmap of the entire inactivating X-chromosome, ordered along the inferred latent time of (b). (**d**) Phase portrait of X-chromosome bin 844 (marked in c). Scatter plot shows the ratio of Dam-LMNB1 (x-axis) and LMNB1 (y-axis) in each cell. Length of arrows reflect the Dam-LMNB1/LMNB1 velocity of bin844 in that cell. Arrows are colored according to differentiation day (left) or latent time (right). (**e**) Same UMAP as in (a), colored according to the ratio between Dam-LMNB1 and LMNB1. (**f**) Same UMAP as in (a), colored according to the Z-score normalized G1 cell cycle-phase enrichment (see Methods). (**g**) Scatter plot of average X Dam-LMNB1 (x-axis) and LMNB1 (y-axis) levels on the Xi. Each dot represents the inactive X allele in one cell, colored according to differentiation day. Distributions above and on the right show average density of day-specific datapoints. Arrow indicates directionality of cells. (**h**) Same scatter plot as in (g), points colored according to the ratio between Z-score normalized G1 and G2 cell cycle-phase enrichment (see Methods).

We next wondered if the loss of genome-lamina interactions of the Xi could be linked to other cellular state characteristics, such as the cell cycle. We determined the cell cycle phase for each single cell using their Hoechst-based DNA content, that was measured during single-cell sorting (Fig. S6e). Interestingly, the precise moment of Xi detachment that we identified through the velocity of genome-lamina interactions (Fig. 6e, Fig. S6f), coincides with a strong enrichment of G1-phase cells (Fig. 6f). Additionally, when projecting cell-cycle phase enrichment over the genome-lamina interaction dynamics of single cells (Fig. 6g-h, Fig. S6g), we observe a striking G2 to G1 phase transition right at the point that the Xi loses its interactions with the nuclear lamina.

Taken together, the Dam&ChIC-inferred chromatin velocity accurately resolves the direction and timing of nuclear localization dynamics of the Xi. In addition, the co-occurrence of detachment from the nuclear lamina and entry into the G1 cell-cycle phase, identify that genome-lamina interactions of the Xi are likely lost upon mitotic exit.

## DISCUSSION

### Advantages and limitations of Dam&ChIC

We developed a multifactorial single-cell chromatin technology, versatile in a twofold manner: 1) for simultaneously profiling two distinct chromatin features at high resolution, and 2) for measuring chromatin-state transitions over time in the same cell. We systematically benchmark the two readouts in Dam&ChIC, by probing diverse combinations of heterochromatic and euchromatic chromatin features. Additionally, when profiling the same chromatin feature – genome-lamina interactions through the nuclear lamina protein LMNB1 – Dam&ChIC detects LADs equally efficiently with either of its two readouts. Overall, we demonstrate that chromatin profiling with Dam&ChIC generates high quality data both in haploid and diploid cells at allelic resolution.

Unlike other methods, Dam&ChIC does not involve *in situ* tagging of the genome. This essentially limits Dam&ChIC to measuring two chromatin features at a time. However, the highly distinct means through which both readouts recover chromatin information, likely contributes to the high-resolution obtained per cell, resulting in unprecedented sensitivity. As a non-Tn5-based approach, Dam&ChIC bypasses the intrinsic bias of other technologies towards more efficient recovery of euchromatic features (Buenrostro et al., 2013; Kaya-Okur et al., 2020), and is therefore also suitable to probe the interplay between features of the constitutive heterochromatin landscape. Despite that some technologies have circumvented the Tn5-intrinsic bias by using Tn5 fusions targeting constitutive heterochromatin (Tedesco et al., 2022) or non-Tn5-dependent genome tagging strategies (Lochs et al., 2023), their sensitivity remains comparatively low. Thus, Dam&ChIC is a suitable approach to examine combinations of chromatin features that range from euchromatic to constitutively heterochromatic, and interrogate their coordinated regulatory function on chromatin, from large domains to individual genes and promoters.

The versatility of Dam&ChIC relies partly on the availability of Dam-POI-expressing cell lines and on high-quality antibodies against targets of interest. A challenge with any DamID-based methodology is the requirement to engineer and integrate the Dam-POI construct for inducible expression in living cells or organisms, which is a significant time investment. However, many cell lines have already been established besides those used in this study, such as for DNA repair proteins (de Luca et al., 2023), and recently the repertoire of DamID has even been extended to single-chain antibodies and engineered chromatin reader domains (Rang et al., 2022). Any of these cell lines are directly compatible with Dam&ChIC, expanding the range of chromatin features that can be profiled. Similar to other antibody-based methods, a high-quality sortChIC readout is highly dependent on the sensitivity and specificity of the antibody used to target the chromatin feature of interest. Profiling of certain chromatin features with very sparse occupancy or transient binding may remain a challenge; nevertheless, a multitude of high-quality antibodies have been characterized and commonly used for epigenomic profiling.

Finally, we note that, as a plate-based approach Dam&ChIC is ideal for medium-throughput experiments, providing flexibility at a low cost compared to high-throughput approaches. Additionally, the plate-based nature of Dam&ChIC offers the opportunity to enrich rare cells of interest based on immunodetection or other cellular properties that can be measured by FACS (Baron et al., 2019). Upscaling the throughput to sample thousands of cells in parallel is challenging and labor-intensive. Nevertheless, we anticipate that future iterations of Dam&ChIC that render it compatible with combinatorial indexing or microfluidic approaches will substantially increase its throughput.

### Recording chromatin transitions over time with Dam&ChIC

Tracking chromatin changes over time in the same cell has been an attractive yet challenging concept. Such technology would enable the study of poorly understood aspects of epigenetic regulation; for example, how past chromatin changes affect present cellular states. With Dam&ChIC we set first steps in these directions, by experimentally integrating the cumulative past chromatin information recorded by scDamID, with snapshot information measured by sortChIC. Previous work identified the dynamic nature of genome-lamina interactions, using a microscopy-based approach (Kind et al., 2013). Consequently, it remains a question whether different types of LADs all undergo a similar degree of dynamics. Here we used the time-resolved Dam&ChIC information to identify extensive variability in genome-lamina dynamics between LADs with different CFs. Based on this premise that Dam&ChIC captures chromatin dynamics, we used the time derivative between both its measurements to infer chromatin velocity. This rationale differs from previous studies that describe chromatin velocity, since those do not directly measure past and present states of the same chromatin feature, but instead are based on the assumption of interdependency between two different simultaneously measured chromatin features in order to extract directional information (Bartosovic & Castelo-Branco, 2022; Tedesco et al., 2022). We use chromatin velocity to study the directional dynamics of LADs and identify diverse LAD behaviors across latent time, such as polarity and focality in the establishment and release of LADs. These observations raise interesting questions on how chromatin is gradually tethered to the nuclear periphery to form LADs (Manzo et al., 2022; Rullens & Kind, 2021).

Currently, Dam&ChIC is limited to recording short-term chromatin dynamics, given the absence of a maintenance machinery in higher eukaryotes that propagates the m6A mark during DNA replication. Consequently, the time window of opportunity to study chromatin dynamics with Dam&ChIC currently spans between two replication cycles. The present Dam&ChIC protocol is therefore very suitable to disentangle the role of chromatin dynamics in cell cycle-related phenomena, or other relatively rapid chromatin changes. We anticipate that future iterations will extend its use over many cellular generations. A candidate to aid propagation of the exogenous m6A mark is the Dam mutant L122A, which has been shown to transfer m6A only on hemi-methylated DNA (Horton et al., 2005), similar to DNMT1 for DNA CpG methylation. In the future, improved engineered versions of such exogenous methyltransferases could make tracking of chromatin dynamics over multiple cell generations feasible.

### Dam&ChIC unravels the order of heterochromatin formation events during XCI

We used Dam&ChIC to study the large-scale epigenetic rewiring that happens during XCI and explore the interplay of chromatin features, with a main focus on the nuclear localization. We provide the first single-cell maps of genome-lamina interactions on the inactivating X chromosome, combined with heterochromatic hPTMs. Previously, microscopy-based studies have described the nuclear periphery or the nucleolus as the preferred nuclear locations of the Xi (Rego et al., 2008; Zhang et al., 2007). More recent findings indicate that the Xi is actively recruited to the nuclear lamina through interactions of Xist with the Lamin B1 receptor (LBR), to enable chromosome-wide silencing (Chen et al., 2016). Surprisingly, our data reveals extensive loss of genome-lamina interactions across mega-base regions on the Xi, that coincide with strong H3K27me3 spreading, a known and early event of XCI (Plath et al., 2003.; Silva et al., 2003; Żylicz et al., 2019). These Xi regions are interspersed with regions that maintain genome-lamina interactions, which coincide with demarcated H3K9me3 domains that pre-exist already before XCI initiation. Although our data seemingly contradicts the observation that the Xi is recruited to the nuclear lamina through interactions of LBR with Xist, these processes may be separated in time, and possibly an initial LBR/Xist-mediated recruitment of Xi at the lamina precedes the dislodgement and H3K27me3 spreading we observe here. In addition, our data is consistent with findings showing that active tethering of the X chromosome to the nuclear lamina does not determine XCI initiation (Pollex & Heard, 2019) and does not exclude that the detaching regions of Xi might instead form interactions with the nucleolus. Interestingly, the H3K9me3 domains that pre-exist already before XCI initiation, demarcate the future Xi regions that resist lamina detachment. This suggests that particularly genome-lamina interactions that are not decorated by H3K9me3, are more vulnerable to detachment.

The observation that precisely those Xi regions that lose genome-lamina interactions strongly accumulate H3K27me3, suggests a potentially antagonistic effect between lamina association and polycomb-deposited H3K27me3 during XCI. A similar phenomenon has recently been described in two unrelated systems, namely K562 cells and preimplantation embryos, where removal of H3K27me3 led to increased genome-lamina interactions (Guerreiro et al., 2023; Siegenfeld et al., 2022). Importantly, our multifactorial analysis implies that the loss of genome-lamina interactions starts just prior to accumulation of H3K27me3. However, both the accumulation of H3K27me3 and the loss of genome-lamina interactions continue to consolidate over XCI. This prompts us to hypothesize that H3K27me3 deposition might further promote exclusion of the Xi from the nuclear lamina, essentially creating a negative feedback loop between genome-lamina interactions and H3K27me3 accumulation. Previous findings have indicated that the accumulation of PRC2-mediated H3K27me3 is preceded by PRC1-mediated H2AK119Ub during XCI, which is enabled only after transcriptional silencing by Xist (Żylicz et al., 2019). Given the close onset of these chromatin events, it is possible that Xist spreading and H2AK119Ub accumulation occur prior to the observed loss in nuclear lamina association and could be involved in the initial release of the Xi from the nuclear periphery. Further studies are necessary to elucidate how the loss of genome-lamina interactions is functionally linked to the spreading of Xist along the X chromosome, transcriptional silencing and the formation of polycomb-mediated heterochromatin.

The time-axis in Dam&ChIC data allowed us to infer chromatin velocity during XCI and resolve the direction of differentiation. More specifically, we aimed to use chromatin velocity during XCI to study the dynamics of genome-lamina interactions in connection to the cell cycle. Our analysis suggests that the loss of genome-lamina interactions occurs directly after mitotic exit, in the G1 phase. Considering additionally our finding that LADs are partially inherited over mitosis, we hypothesize that the mitotic exit itself might offer a window of opportunity for detachment to take place, since the chromosomes need to find their way back to the re-assembled nuclear lamina. While events that prime detachment of Xi from the nuclear lamina may potentially occur prior to mitosis, the detachment might only materialize after mitotic exit. Future studies can uncover what triggers release of the Xi from the nuclear lamina, and how this phenomenon coordinates with mitotic exit. Taken together, our comprehensive single-cell analysis highlights the temporal dynamics of spatial positioning of the X chromosome and provides new insights into heterochromatin formation during XCI.

### Conclusion

Several aspects of chromatin regulation have been poorly understood due to the limitations posed by conventional technologies. Dam&ChIC introduces unique possibilities to investigate in depth the interplay between diverse chromatin factors, and to track retrospectively temporal changes within the same cell. Our single-cell multifactorial chromatin data and chromatin velocity implementations elucidate aspects of chromatin regulation, that are involved in the inheritance of chromatin upon mitosis and the formation of heterochromatin during XCI. These properties establish Dam&ChIC as a unique tool in the rapidly growing set of single-cell chromatin technologies.

## ACKNOWLEDGEMENTS

We thank members of the Kind and van Oudenaarden labs for useful feedback, discussions throughout the project and critical reading of the manuscript. We thank Reinier van der Linden and Anita Pfauth (Hubrecht FACS facility) for performing single-cell sorting. We thank Koos Rooijers for developing data processing methods. This work was supported by ERC Consolidator grant (ERC-CoG 1010002885-FateID) and a Nederlandse Organisatie voor Wetenschappelijk Onderzoek (NWO) VIDI grant (016.161.339) to J.K. P.Z. was funded by SNF (P2BSP3-174991), HFSP (LT000209/2018-L) and Marie Skłodowska-Curie Actions (798573). The work in the lab of A.v.O. is supported by (ERC-AdG 101053581-scTranslatomics), and Nederlandse Organisatie voor Wetenschappelijk Onderzoek (NWO) TOP award (NWO-CW714.016.001). The Oncode Institute is partially funded by the KWF Dutch Cancer Society. We thank the Utrecht Sequencing Facility (USEQ) for providing sequencing service and data. USEQ is subsidized by the University Medical Center Utrecht and The Netherlands X-omics Initiative (NWO project 184.034.019).

## AUTHOR CONTRIBUTIONS

S.K., P.Z. and J.K. conceived the method. S.K., P.M.J.R., P.Z. and J.K designed the project, with input from K.L.d.L. S.K. and P.Z. developed the method. S.K. performed all experiments. P.M.J.R performed all computational analyses and developed software. S.K. and P.M.J.R. curated and validated the data. P.Z., K.L.d.L. and S.S.d.V. assisted with experiments. T.K. and P.Z. performed preliminary analyses. P.M.J.R. and S.K. visualized the data. J.K., P.Z. and A.v.O. supervised the project. J.K and A.v.O. acquired funding. S.K., P.M.J.R., P.Z. and J.K. wrote the manuscript.

## METHODS

### Experimental Methods

#### Dam-POI cell lines

KBM7 cells expressing Dam-LMNB1 or untethered Dam under the control of the destabilizing domain (DD) system were generated previously using lentiviral transductions (Kind et al., 2015). Mouse F1 hybrid Cast/EiJ x 129/Sv embryonic stem cells (ESCs) were a kind gift from the Joost Gribnau laboratory. These mESC lines are derived from female cells and do not contain a Y chromosome. The Dam-LMNB1 or Dam-scFv-H3K27me3 constructs were previously integrated in these cells using CRISPR targeting (Guerreiro et al., 2023; Rang et al., 2022). Both ESC lines express the Dam-POI under the control of the AID degron system. In Dam-scFv-H3K27me3 ESCs, expression of the Dam fusion protein is additionally controlled by tamoxifen-induced nuclear translocation via the estrogen receptor (ER).

#### Cell culture

All cell lines were cultured in a humidified chamber at 37°C in 5% CO2 and tested for mycoplasma at a regular basis. Human haploid KBM7 cells were cultured in suspension in IMDM medium (Gibco, 12440053) supplemented with 10% FBS (Sigma, F7514) and 1% Pen/Strep (Gibco, 15140122). Mouse F1 hybrid ESCs were cultured on a monolayer of irradiated primary mouse embryonic fibroblasts (MEFs) in CM+/+, as described previously (Rang et al. 2022). CM+/+ consists of G-MEM (Gibco, 11710035) supplemented with 10% FBS, 1% Pen/Strep, 1× GlutaMAX (Gibco, 35050061), 1× non-essential amino acids (Gibco, 11140050), 1× sodium pyruvate (Gibco, 11360070), 0.1 mM β-mercaptoethanol (Sigma, M3148) and 1000 U/ml ESGROmLIF (EMD Millipore, ESG1107). For passaging, mouse ESCs cells were washed once with PBS0, treated with TrypLE (Gibco, 12605010) for 4 min at 37oC, neutralized with serum-containing medium, and after one wash they were plated at the desired density. Prior to differentiation, mouse ESCs were passaged at least once in feeder-free conditions, on 0.1% gelatin-coated 6-wells, and cultured in medium containing 40% CM +/+ and 60% BRL-conditioned CM +/+ medium. Expression of the Dam-POI in the mouse ESC lines was suppressed by addition of 0.5 mM indole-3-acetic acid (IAA; Sigma, I5148).

#### Induction of Dam-POI expression

In KBM7 cells, Dam-LMNB1 and the untethered Dam were induced with the addition of 0.5 nM Shield-1 (Glixx Laboratories Inc, GLXC-02939) that enables stabilization of the protein. In hybrid mESCs, 24 hours prior to induction the cells were cultured in 1 mM indole-3-acetic acid (IAA; Sigma, I5148), to ensure protein degradation. For induction the cells were washed three times with PBS and fresh medium without IAA was added, followed by the addition of 1 uM 4-Hydroxytamoxifen (4-OHT; Sigma, SML1666), when required. In detail, Dam-LMNB1 mESCs were induced for 6 hours without IAA and Dam-scFv-H3K27me3 mESCs were induced for 20 hours without IAA and 6 hours with 4-OHT.

#### *In vitro* differentiation of mouse hybrid ESCs with Ascorbic Acid (Vitamin C)

Dam-POI expressing mouse hybrid ESCs were passaged at least once in feeder-free conditions and cultured as described above. The culture was depleted from the remaining MEFs and 24-48 hours later confluent wells were passaged and plated in AA differentiation medium, defined as IMDM supplemented with 15% FBS, 1% Pen/ Strep, 1X GlutaMAX, 1X non-essential amino acids, 50 μg/ml Ascorbic Acid (Sigma, A4544) and 0.1 mM β-mercaptoethanol (Loda et al., 2017; Robert-Finestra et al., 2021). Expression of the Dam-POI was suppressed by addition of 0.5 mM IAA. The medium was refreshed every day during the course of differentiation.

#### Cell-cycle synchronization with double thymidine block

Confluent cultures of KBM7 cells were passaged 24 hours prior to the initiation of synchronizations and 24 hours later received 2 mM thymidine (Sigma, T1895) for 15 hours, followed by 9 hours of release, followed by a second thymidine block for 13 hours. Synchronization tests were performed prior to Dam&ChIC experiments, in order to define the cell-cycle progression of these cell lines based on the DNA content at 0, 4, 6, 7, 8, 9, 10, 11, 12, 13 and 14 hours after release from the second thymidine block. Cells were harvested at the desired time points, counted on a cytometer, and after one wash with PBS0 0.5 - 1 million cells were fixed with 70% ethanol at -20°C overnight. Afterwards, the fixed cell suspension was washed once with ice-cold PBS0 and stained in PBS0 containing 2.5 ug/ml Hoechst 34580 (Sigma) for 48 hours at 4°C. To standardize the stainings, equal cell numbers were used from each time point. DNA content measurements were performed on a BD LSR Fortessa FACS, and subsequent analyses were done using FlowJo. To follow KBM7 cells over mitosis with Dam&ChIC, KBM7 cells were synchronized with a double thymidine block as above. During the second thymidine block, 0.5 nM Shield-1 was added to enable stabilization of Dam-LMNB1. Afterwards, both thymidine and Shield-1 were washed out, and cells were released to progress through S-phase. Cells in G2 were harvested 8 hours post-release and cells after mitotic exit in early G1 were harvested 13 hours post-release.

### Preparation of samples for Dam&ChIC

#### Cell harvesting and permeabilization

Nuclei isolation was done like described before by Zeller et al., 2023. In short, cells were collected in 15 ml falcon tubes and washed three times with room temperature PBS0. After counting, 0.5-1 million cells per staining were used for permeabilization in ice-cold Wash Buffer 1 (20mM HEPES pH 7.5, 150mM NaCl, 66.6 ug/ml Spermidine (Sigma, S2626-1G), 1x cOmplete protease inhibitor cocktail (Roche, 11836170001), 0.05% Saponin (Sigma, 47036-50G-F), 2mM EDTA) and washed once at 300g for 4 minutes at 4°C, before addition of the primary antibody. Subsequent washes were done with the same centrifuge settings.

#### Cell harvesting, ethanol fixation and permeabilization

For ethanol fixation, cells were collected in 15 ml falcon tubes and washed three times with room temperature PBS0. After counting, 0.5-1 million cells were fixed with 70% ice-cold ethanol for 1 hour at -20°C. Fixed cells were washed once and permeabilized in ice-cold Wash Buffer 1F (20mM HEPES pH 7.5, 150mM NaCl, 1x cOmplete protease inhibitor cocktail, 0.05% Tween). Afterwards, samples were stained with CellTrace dyes (see below) or frozen at -80°C for long-term storage in Wash buffer 1F that contains additionally 66.6 ug/ml Spermidine, 2mM EDTA and 10% DMSO.

#### Staining with CellTrace dyes

Upon fixation and washing, 0.5-1 million cells were stained in 1 ml Wash buffer 1F containing 0.25 ul of a CellTrace dye (CellTrace CFSE, CellTrace Far Red, CellTrace Yellow) at 4° C for 20 minutes protected from light, as described before (Yeung et al., 2023). When multiple dyes were used per sample, each was used equimolarly. The stainings were quenched with the addition of 50 ul Rat Serum (Sigma, R9759) and incubation at 4°C for 10 minutes protected from light. Samples were washed once with Wash buffer 1F that contains additionally 66.6 ug/ml Spermidine and 2mM EDTA, and resuspended in the same buffer for subsequent antibody stainings. Part of the sample was aliquoted and frozen at -80°C for long term storage, as described above.

#### Multiplexing of samples based on CellTrace stainings

Samples stained with different combinations of CellTrace dyes were mixed in one “super-sample” to enable parallel processing and thereby minimize batch effects, as described before (Yeung et al., 2023). Specifically, samples from each day of the Vitamin C differentiation time-course were stained with a unique combination of CellTrace dyes, including CellTrace CFSE, CellTrace Far Red and CellTrace Yellow or left unstained. This enabled the distinction of up to eight populations in the “super-sample” based on fluorescence. In the synchronization and mitosis experiment, the G2 sample was stained with CellTrace Yellow and CellTrace CFSE, and the early G1 sample was stained with CellTrace CFSE, CellTrace Far Red and CellTrace Yellow. In the latter experiment, more CellTrace-stained control samples were mixed in the same “super-sample”, but only the relevant stainings are presented here for simplicity (Fig. S3b). For all experiments, equal cell numbers from each sample were combined together to a final total amount of 0.5-1 million cells for subsequent antibody staining.

#### Antibody staining and pA-MNase tethering

Nuclei or fixed cells were stained with the primary antibody overnight at 4°C on a roller. The concentrations used varied per antibody and LOT number, and were defined by titrations done in bulk samples prior to single-cell experiments. Specifically, anti-LMNB1 (Abcam, ab16048) was used at 1:200 or 1:400; anti-H3K4me1 (Abcam, ab8895) at 1:400; anti-H3K27me3 (Cell Signaling Technologies, 9733S) at 1:200; anti-H3K9me3 RM389 (Thermofisher, MA5-33395) at 1:200; anti-H3K4me3 (Thermofisher, MA5-11199) at 1:400. After one wash with Wash Buffer 2 (for nuclei; 20mM HEPES pH 7.5, 150mM NaCl, 66.6 ug/ml Spermidine, 1x cOmplete protease inhibitor cocktail, 0.05% Saponin) or Wash Buffer 2F (for fixed/permeabilized cells; 20mM HEPES pH 7.5, 150mM NaCl, 66.6 ug/ml Spermidine, 1x cOmplete protease inhibitor cocktail, 0.05% Tween), pA-MNase was added at a final concentration of 3 ng/ul for KBM7 samples and 0.6 ng/ul for ESC and Vitamin C samples, in the respective buffers. Hoechst 34580 was added in the same mix at a final concentration of 2.5 ug/ml. The samples were incubated in a roller at 4°C for 1 hour, followed by two washes in the same wash buffer, and finally passed through a cell-strainer for sorting.

#### Single-cell sorting and FACS gating strategies

FACS sorting was performed on the BD FACS Influx Cell Sorter System or BD FACS Jazz Cell Sorter System, with sample and plate cooling. Gating was based on size (forward-side scatter), exclusion of doublets, and Hoechst-defined DNA content (Fig. S3b). For experiments in which different samples were multiplexed, for example in the mitosis and XCI experiments, an additional gating strategy based on fluorescence levels of the CellTrace dyes ensured distinction of the different populations (Fig. S3b). In the mitosis experiment, G1 and G2 cells were additionally selected as positive for CellTrace CFSE and CellTrace Yellow (CTY), and were sorted on a combination of Hoechst and CellTrace Far Red (CTFR) to ensure selection of pure synchronized cells in G2 and G1 (Fig. S3b). One cell per well was sorted in hard-shell 384 well-plates that were pre-filled with 5 uL/well mineral oil using the Freedom EVO liquid handling platform (Tecan), and 100 nL/well Wash buffer 3 (for nuclei; 20mM HEPES pH 7.5, 150mM NaCl, 66.6 ug/ml Spermidine, 0.05% Saponin) or Wash buffer 3F (for fixed/permeabilized cells; 20mM HEPES pH 7.5, 150mM NaCl, 66.6 ug/ml Spermidine, 0.05% Tween). After sorting, plates were, sealed with aluminum seals (Greinier, 676090), spun at 2000g for 1-2 minutes at 4°C, and kept at 4°C until further processing.

#### Single-cell Dam&ChIC

Liquid dispensions were performed using the Nanodrop II liquid handler (Innovadyne) and adapter dispensions were performed using the Mosquito LV (STP Labtech), as described before (Markodimitraki et al., 2020; Rang et al., 2022; Zeller et al., 2023). In short, pA-MNase in single nuclei was activated after dispension of 100 nL per well of Activation Solution (Wash buffer 3 or Wash buffer 3F, containing 4mM CaCl2) to a total volume of 200 nL per well. The plates were incubated at 4°C for exactly 30 minutes, followed by dispension of 200 nL per well of Stop Solution (0.04M EGTA, 1.5% NP40, 2 ug/ul Proteinase K) to a total volume of 400 nL. For lysis and proteinase K treatment, plates were incubated at 65°C for 6 hours, followed by heat inactivation at 80°C for 10 minutes. Plates were stored at -20°C until further processing. 200 nL per well of Blunt-ending mix (0.02 U Klenow, 0.04 U T4 PNK, 0.5mM dNTPs, 3mM ATP, 2.5mM MgCl2, 2.5% PEG8000, 0.6 ug/ul BSA, PNK buffer) were dispensed to a total volume of 600 nL, followed by incubation at 37°C for 30 minutes and heat inactivation at 75oC for 20 minutes. Next, digestion with DpnI mix (0.2 U DpnI, PNK buffer, 0.5 ug/ul BSA) was done with dispension of 400 nL to a total volume of 1000 nL, followed by incubation at 37°C for 8 hours and heat inactivation at 80°C for 20 minutes. Finally, the content of each well was barcoded through dispension of 50 nL of DamID2 adapters at a final concentration of 25 nM and dispension of 950 nL Ligation mix (0.375 U T4 Ligase, Ligase buffer) to a final volume of 2000 nL, followed by incubation at 16°C for 16 hours and heat inactivation at 65°C for 10 minutes. After each dispension step, the plates were sealed with aluminum plate seals.

#### Library preparation

The content of all wells in a plate was pooled by centrifugation of the 384-well plates on collection plates, transferred to eppendorfs, and the aqueous phase was separated from the oil phase by a few rounds of centrifugation and transfers to clean tubes. Sample purification with beads followed, like described before (Hashimshony et al., 2016), by incubation of the sample with 0.8 volume 1:10 diluted beads (CleanNGS; GC Biotech, CNGS-0050) in bead-binding buffer, two washes with 80% Ethanol and elusion with 7 uL ultra-pure nuclease-free water. The eluted sample was then in vitro transcribed at 37°C for 14 hours, using the MEGAScript T7 transcription kit (Invitrogen, AMB13345). The amplified RNA (aRNA) was purified using 0.8 volume undiluted beads, washed three times with 80% Ethanol, and eluted in 13 uL ultra-pure nuclease-free water. The quality of the eluate was evaluated using the Agilent RNA 6000 Pico Assay, followed by library preparation, like described previously (Hashimshony et al., 2016), with a few adjustments. In particular, a maximum of 6 uL of 100-200 ng aRNA was used for reverse transcription, followed by 8-11 cycles of library PCR, depending on the amount of input aRNA, and the amplified material was purified twice, as described above, before final elusion in 13 uL of ultra-pure nuclease-free water. Libraries were quantified using the Qubit sdDNA High Sensitivity Assay and the Agilent High Sensitivity DNA Assay, and subsequently sequenced with single-end or paired-end sequencing on the Illumina NextSeq500 (75-bp reads) or the Illumina NextSeq2000 (100-bp reads).

### Data processing

#### Processing Dam&ChIC data

Dam&ChIC data processing is largely based on the workflow and scripts described in Rooijers et al., 2019 and Markodimitraki et al., 2020, but adapted to allow for computational separation of scDamID and sortChIC derived reads and further processing of sortChIC reads. The key steps of the procedure are described below (detailed explanation and code are available at https://github.com/KindLab/DamChIC).

##### Raw data pre-processing

All reads are demultiplexed by comparing their sample barcode (e.g. single-cell barcode) located at the start of each read (R1 in case of paired-end sequenced libraries) conform to the read layout of 5′-[3 nt UMI][8 nt barcode] TC[gDNA]-3′ for scDamID and 5′-[3 nt UMI][8 nt barcode]WN[gDNA]-3′ for sortChIC reads, to a reference list of barcodes and zero mismatches between the observed barcode and reference are allowed. Sample barcodes and UMIs are trimmed off the reads to only retain gDNA sequences. The UMI sequences were appended to the read name for downstream processing. Per sample barcode, scDamID and sortChIC reads remain mixed at this point.

##### Sequence alignments

After preprocessing, the reads were aligned to a reference genome using bowtie2 (v. 2.4.1) with parameters: ‘‘--seed 42 --very-sensitive -N 1’’. Before alignment a ‘GA’ dinucleotide was prepended to the reads that has effectively been digested off scDamID fragments during DpnI gDNA digestion. For human samples, reads were aligned to reference genome hg19 (GRCh37) and mouse samples to mm10 (GRCm38). For allele-specific processing for X-chromosome inactivation experiments, data were aligned to reference genomes generated by imputing 129/Sv and CAST/ EiJ single nucleotide polymorphisms obtained from the Sanger Mouse Genomes project (Keane et al., 2011) onto the mm10 reference genome. The reads were assigned to either genotype by aligning reads to both references. Reads that aligned with lower edit distance (SAM tag ‘NM’) or higher alignment score (SAM tag ‘AS’) in case of equal edit distance to one of the genotypes were assigned to that genotype. Reads aligning with equal edit distance and alignment score to both genotypes were considered of ‘ambiguous’ genotype. Reads that yielded an alignment with mapping quality (BAM field ‘MAPQ’) lower than 10 were discarded.

##### In silico separation of scDamID and sortChIC reads

Following read alignment with ‘GA’ prepended dinucleotide, reads were considered scDamID in case of perfect alignment on genomic GATC motifs. Residual reads were considered sortChIC and their original sequences (i.e., without prepended GA dinucleotide) were realigned using hisat2 (v. 2.1.0) with parameters: ‘‘-- seed 42 --no-spliced-alignment --mp ‘2,0’ --sp ‘4,0’”. These sortChIC-considered reads were further pruned given MNase’s strong adenine or thymidine-nucleotide cut preference (Fig. S1c); only sortChIC reads were kept that start with an ‘A’ or ‘T’ nucleotide. In addition, we observed that a larger fraction of sortChIC reads (i.e., that do not map on a genomic GATC motif) are ‘TC’-dinucleotide starting in Dam&ChIC libraries than in sortChIC libraries (Fig. S1d) and hence decided to exclude TC-starting reads from Dam&ChIC-derived sortChIC libraries.

##### PCR duplicate filtering

To exclude PCR read-duplicates, both scDamID and sortChIC read counts were collapsed based on genomic position, strand and unique molecular identifier (UMI) sequence. In case of haploid KBM7 cells or allele-specific data processing, multiple reads with the same UMI count as 1, for diploid data processing as 2. The number of observed unique UMIs was taken as the number of unique methylation events for scDamID or unique MNase cut sites for sortChIC.

##### Filtering of samples

To exclude single-cell samples from the analysis that failed we applied construct-specific cutoffs on UMI-unique reads for scDamID: 1,000 for Dam and Dam-scFv-H3K27me3 (Rang et al., 2022), and 5,000 for Dam-LMNB1. A general cutoff of 1,000 UMI-unique reads per sample was applied for ChIC. Samples in allele-specific analysis were included if ≤200 UMI-unique reads could be assigned to both parental alleles for scDamID as well as sortChIC-derived reads.

#### ChIP-seq data processing

External K562 ChIP-seq datasets were downloaded from the ENCODE database (Davis et al., 2018). ChIP-seq datasets from K562 cells used for this manuscript include H3K4me3 (ENCSR668LDD), H3K36me3 (ENCSR000DWB), H3K27me3 (ENCSR000EWB) and H3K9me3 (ENCSR000APE). ChIP-seq reads were aligned using bowtie2 (v. 2.4.1) with parameters: ‘‘-- seed 42 --very-sensitive -N 1’’. ChIP-seq data were binned using the “bamCoverage” function from DeepTools (v. 3.3.2) with parameters: ‘‘--ignoreDuplicates -- minMappingQuality 10’’. NarrowPeak calls were likewise downloaded from the ENCODE database from the corresponding ChIP-seq datasets.

### Downstream Analyses

#### Binning and calculation of OE values

For most analyses, scDamID and sortChIC data were binned using consecutive non-overlapping 100-Kb bins. For gene-based analysis (Fig. 1c,d) data were binned at a higher resolution of 1-Kb. To be able to calculate OE values, an “expected” vector for scDamID data was generated in silico by generating sequences of 65 nt (in both orientations) from the reference genome and aligning and processing them identically to the data. By binning the in silico generated reads, the maximum amount of mappable unique events per bin was determined (Fig. S1g). The expected for sortChIC was generated experimentally using MNase-H3 (Fig. S1g), unless stated otherwise. OE values were calculated as described in Rooijers et al., 2019.

#### Data binarization, Contact Frequency and combined single-cell heatmap

If specifically stated, data was binarized by setting a threshold of log2(≥1) based on the bimodal distribution of OE values of 100-Kb genomic bins (Fig. S2a) (Kind et al., 2015). The contact frequency (CF) metric is used throughout the manuscript and is defined by calculating the fraction of samples (single cells) that meet the binarization threshold for a given 100-Kb bin (Kind et al., 2015). The combined single-cell heatmap presented in Fig. 3c is determined by first binarizing the 100-Kb binned data of both Dam&ChIC constituents separately, resulting in two presence-absence *m* × *n* matrices; 1) scDamID defined as

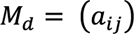

and 2) sortChIC defined as

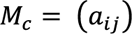

where *i* is a cell and *j* a100-Kb bin.Both matrices are combined in one by simple summation as

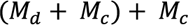

In the resulting combined matrix 0 denotes absence of both, 1 presence only of scDamID, 2 presence only of sortChIC and 3 presence of both.

#### Dimensionality reduction on KBM7 Dam&ChIC data

The UMAP presented in Fig. 1f was computed by first calculating the first 50 principal components (Python; sklearn v. 1.0.2) on the RPKM normalized single-cell data binned at 100-Kb resolution, on which UMAP (Python; umap v. 0.5.2) was then run.

#### (single-cell) LAD calling

Bulk LADs presented in Fig. S1c are called by setting a threshold of log2(≥1) based on the bimodal distribution of OE values of 100-Kb genomic bins Fig. S1a (Kind et al., 2015). Notably, LAD calls <300-Kb were excluded and LAD calls intervened by ≤100-Kb were combined into one. For single-cell LAD calling used for Fig. 2, LADs we first separately called on pseudobulk profiles of both Dam&ChIC readouts as described above. Only LADs that were identified by both readouts were kept for single-cell analysis (Fig. S2c). The resulting LAD coordinates were in each cell tested for having an average OE log2(≥1), if this requirement was met, the LAD was called in the given cell. This resulted in two LAD-based binary presence-absence matrices; 1) scDamID defined as

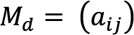

and 2) sortChIC defined as

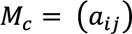

where *i* is a cell and *j* a LAD.

#### Jaccard Z-score normalization by matrix permutation

To compare the single-cell co-ocurrence of scDamID and sortChIC identified LADs between different Dam&ChIC experiments and across LADs with different CF’s, the Jaccard similarity was index was computed, defined as

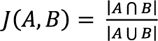

where *A* represents the scDamID and *B* the sortChIC measurement from the same cell (Fig. S2f, Fig. 3b, Fig. S3e). Two inherent problems are posed on this analysis that might introduce undesirable bias and ultimately unreliable comparison between different experiments or LADs: 1) typical sparsity of single-cell data results in non-uniformly distributed signal dropout and 2) binary similarity metrics can be sensitive to site prominence that varies between different LADs, but which does not reflect true co-ocurrence of scDamID and sortChIC signal. To solve both problems, we applied a previously described matrix randomization algorithm (Strona et al., 2014), to permute the abovementioned presence-absence LAD matrix *Mc n*-times (*n* = 100), without altering row and column totals. The resulting randomized matrices were used to compute Jaccard similarity with LAD matrix *M*d to serve for Z-score normalization, which can be written as

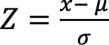

where *x* is the similarity score of the observed, μ the mean similarity and σ the standard deviation of the random controls.

#### Chromatin velocity

Chromatin velocity analysis presented in Fig. 2 and 6 were performed using the scVelo Python package v.0.2.5 (Bergen et al., 2020). An average OE value was calculated in the abovementioned LAD calls for each cell for both the Dam-LMNB1 and LMNB1 Dam&ChIC readout. The resulting Dam-LMNB1 matrix was provided to scVelo as the “spliced” layer and the LMNB1 matrix as “unspliced”.

#### XCI dimensionality reduction and trajectory inference

Dimensionality reduction of the XCI analysis was performed with Seurat v.4.1.0 (Hao et al., 2021) and Signac v.1.3.0 (Stuart et al., 2021). The UMAP introduced in Fig. 5b is based on H3K27me3 signal on autosomal genes (i.e., 5-Kb upstream of TSS + gene body). Different batches (i.e., Dam&ChIC experiments) were integrated using Harmony v.0.1.0 (Korsunsky et al., 2019). Pseudotime inference was performed using Monole3 v.1.0.0 (Cao et al., 2019).

#### Z-score normalized cell cycle analysis on XCI data

The cell cycle analysis on the XCI dataset presented in Fig. S6 is based on Hoechst staining of the DNA of cells before they are sorted in plates, which reflects DNA content and therefore enables to categorize cells into the different cell cycle phases (i.e., G1, S or G2). However, these phases are not equally divided across single cells, as the majority of the cells in culture are in G1-phase (Fig. S6e,g). This poses a problem on the cell cycle analysis presented in Fig. 6f,h, that may lead to overestimation of one stage over another. To solve this, we performed Z-score normalization of the observed cell cycle distributions within k-nearest neighbors (*kNN*, *k* = 100). For each neighborhood, the observed cell cycle distribution was compared to *n*-times randomly sampled controls (*n* = 100). Z-score normalization was defined as follows

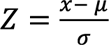

where *x* is the cell cycle distribution of the observed, μ the mean similarity and σ the standard deviation of the randomly sampled controls.

##### Data and Code availability

All genomic data generated in this study was deposited on the NCBI Gene Expression Omnibus (GEO) portal and is publicly available as of the date of publication under accession number GSE247458. Code is available at https://github. com/KindLab/DamChIC. Additional information required to reanalyze data reported in this manuscript is available by the Lead Contact upon request.

---

**Figure S1.**
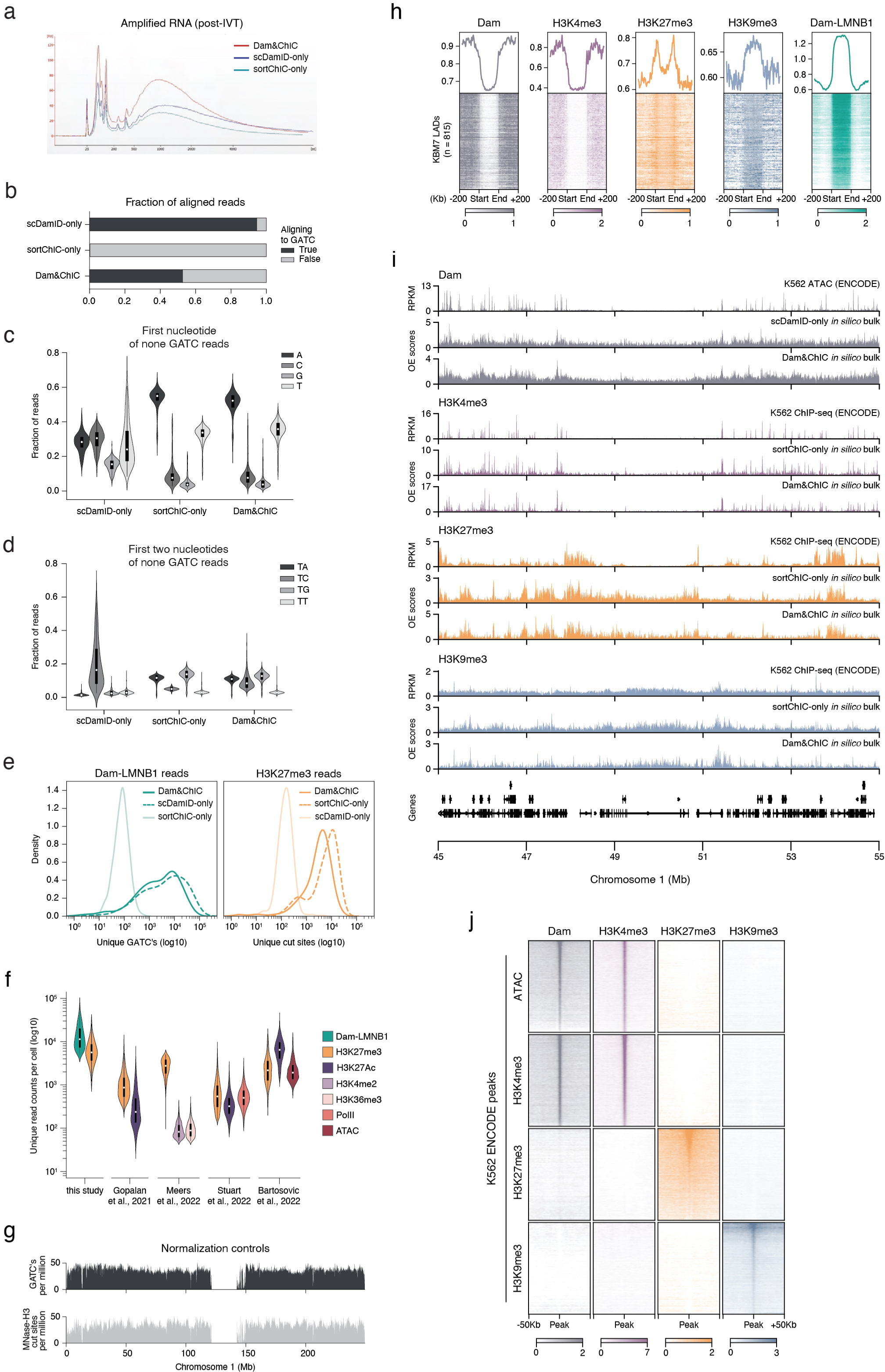
(a) Bioanalyzer plot showing the amount and size distribution of amplified RNA (aRNA) samples, derived after in vitro transcription (see Methods) with the Dam&ChIC protocol. Dam&ChIC was performed to profile Dam-LMNB1 and H3K27me3, alongside respective control scDamID-only and sortChIC-only experiments. Each aRNA product was derived from one 384-well plate containing 380 cells, pooled together prior to amplification. The samples were processed in parallel. (**b**) Comparison of the fraction of reads mapping on a genomic GATC motif (black) between Dam&ChIC, scDamID-only and sortChIC-only. (c) Violin plots showing a quantification of the first nucleotide of invalid DamID reads (i.e., not mapping on genomic GATC motifs) for Dam&ChIC, scDamID-only and sortChIC-only. (**d**) Violin plot depicting a quantification of the first two nucleotides of invalid DamID reads for Dam&ChIC, scDamID-only and sortChIC-only. (**e**) Distribution of the unique number of Dam-LMNB1 (left) or H3K27me3 (right) reads recovered per cell by scDamID-only, sortChIC-only or Dam&ChIC. (**f**) Comparison of unique number of reads obtained per cell between Dam&ChIC and other recently published multifactorial chromatin profiling methods (Gopalan et al., 2021; Meers et al., 2022; Bartosovic et al., 2022; Stuart et al., 2022). (**g**) Genomic distribution of in silico GATC mappability (see Methods) (top, black) and experimental pA-MNase H3 mappability (bottom, gray). (**h**) Heatmaps showing chromatin features profiled with Dam&ChIC aligned on LADs (n = 815) defined in KBM7 cells (Rooijers et al., 2019). Profiles above show the scaled averages of the heatmaps for each mapped chromatin feature. (**i**) Genomic tracks of average (in silico bulk) Dam&ChIC profiles compared to corresponding sortChIC-only datasets in KBM7 cells and corresponding ENCODE (ENCODE 2012) ChIP-seq or ATAC-seq datasets in K562 cells. (**j**) Tornado plots depicting pseudobulk Dam&ChIC data aligned on ENCODE ChIP-seq or ATAC-seq peak coordinates in K562 cells. Peaks (rows) are ordered on the peak signal of corresponding Dam&ChIC dataset (i.e., ATAC-seq peaks are ordered on Dam scDamID data, H3K4me3 peaks are ordered on H3K4me3 sortChIC data and so on).

**Figure S2.**
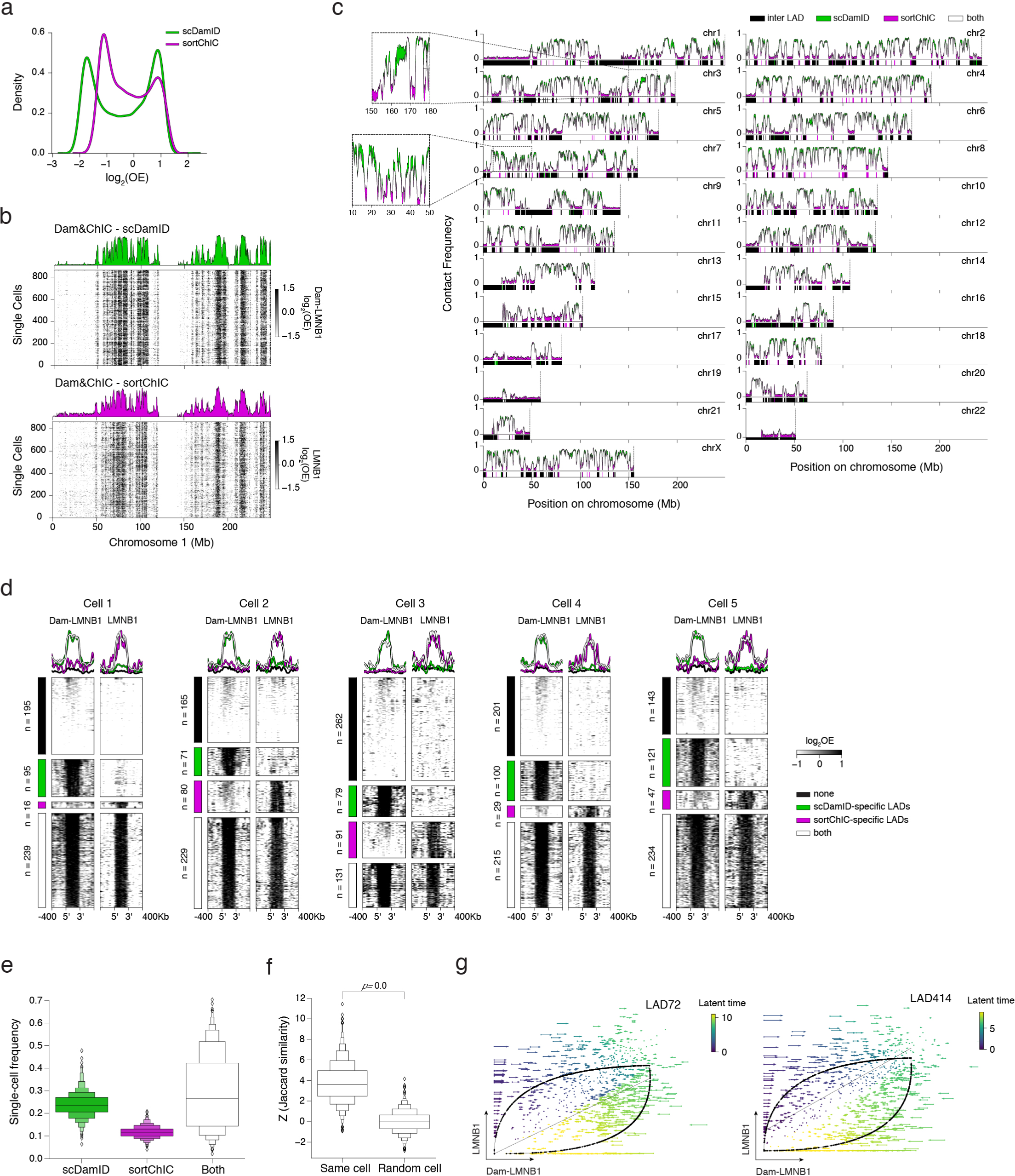
(**a**) Distribution of log-transformed observed over expected (OE) values of 100Kb binned pseudobulk Dam&ChIC data. (**b**) Single-cell heatmaps showing log-transformed observed over expected (OE) LAD signal (see Methods) in 100Kb genomic bins on chromosome 1 of either the scDamID (top) or the sortChIC (bottom) readout. (**c**) Comparison of LAD contact frequency (see Methods) on depicted chromosomes between the scDamID (green) and sortChIC (magenta) measurement of Dam&ChIC. Overlapping signal is shown in white. LAD calls (see Methods) are shown below each track. (**d**) Five example single cells aligned and grouped for all bulk-based LADs according to their detection in the given cell by neither measurement (black), scDamID (green), sortChIC (magenta) or both (white). scDamID (left) and sortChIC (right) measurements are split, and respective enrichment for each category is shown as line plots at the top. LAD signal is shown as log-transformed observed over expected (OE) in 100Kb genomic bins. (**e**) Quantification of the frequency by which each LAD is measured in a single cell by scDamID, sortChIC or both. (**f**) Z-score normalized Jaccard similarity (see Methods) between the scDamID and sortChIC readouts from the same cell and any two random cells. (**g**) Example phase portraits for LAD72 and LAD414, colored according to their latent time, like in (3f).

**Figure S3.**
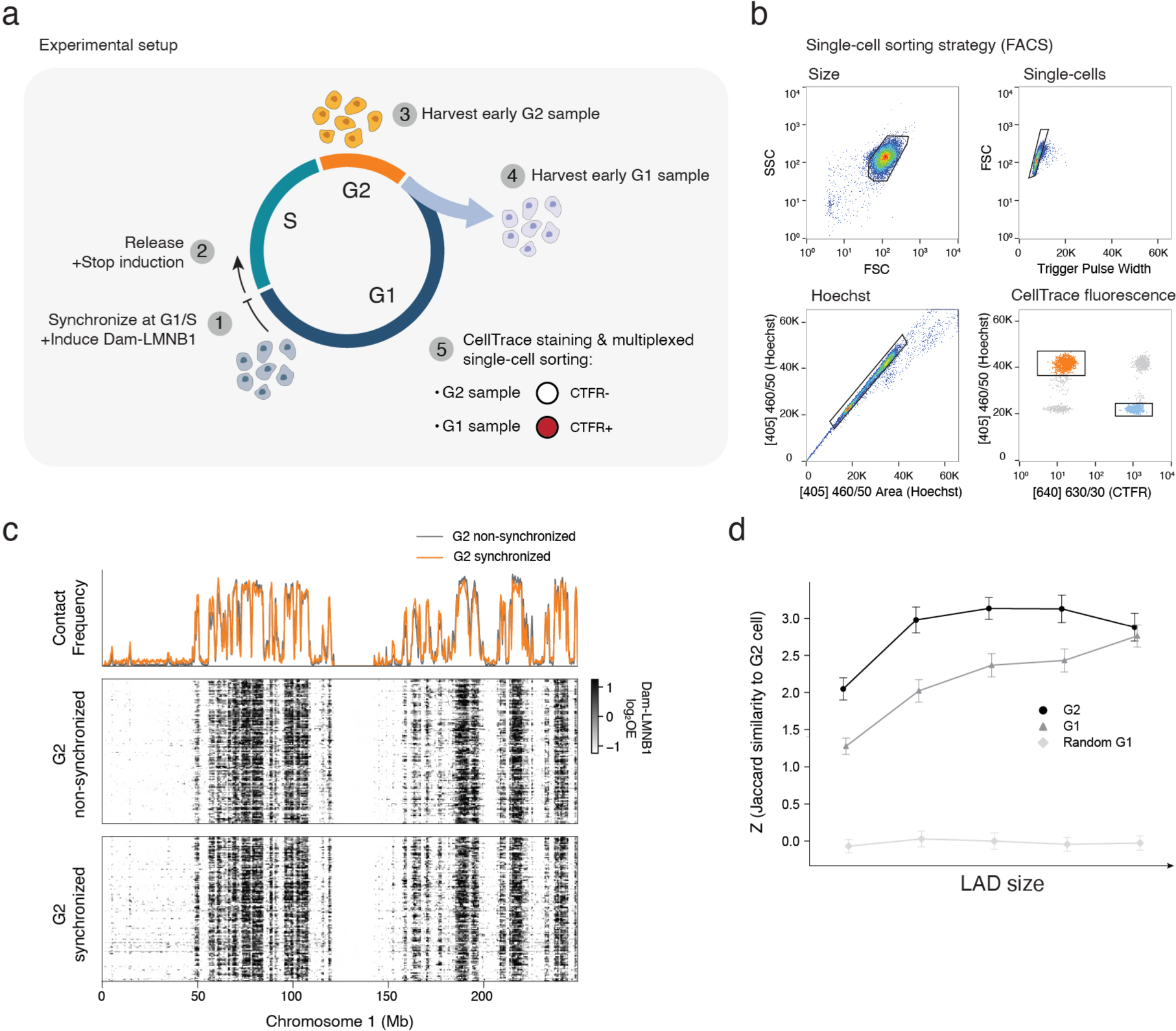
(**a**) Schematic of the experimental setup for the mitotic inheritance Dam&ChIC dataset (see Methods). (**b**) Single-cell sorting strategy (see Methods), including the used gates based on size, singlets, Hoechst staining and CellTrace stainings. CellTrace Far Red is noted as CTFR. (**c**) Single-cell heatmaps and corresponding Contact Frequency quantifications, comparing G2 synchronized with G2 non-synchronized cells. (**d**) Z-score normalized Jaccard similarity between scDamID and sortChIC readouts within G2 cells, within G1 cells, and between random G1 cells, across different LAD size quantiles.

**Figure S4.**
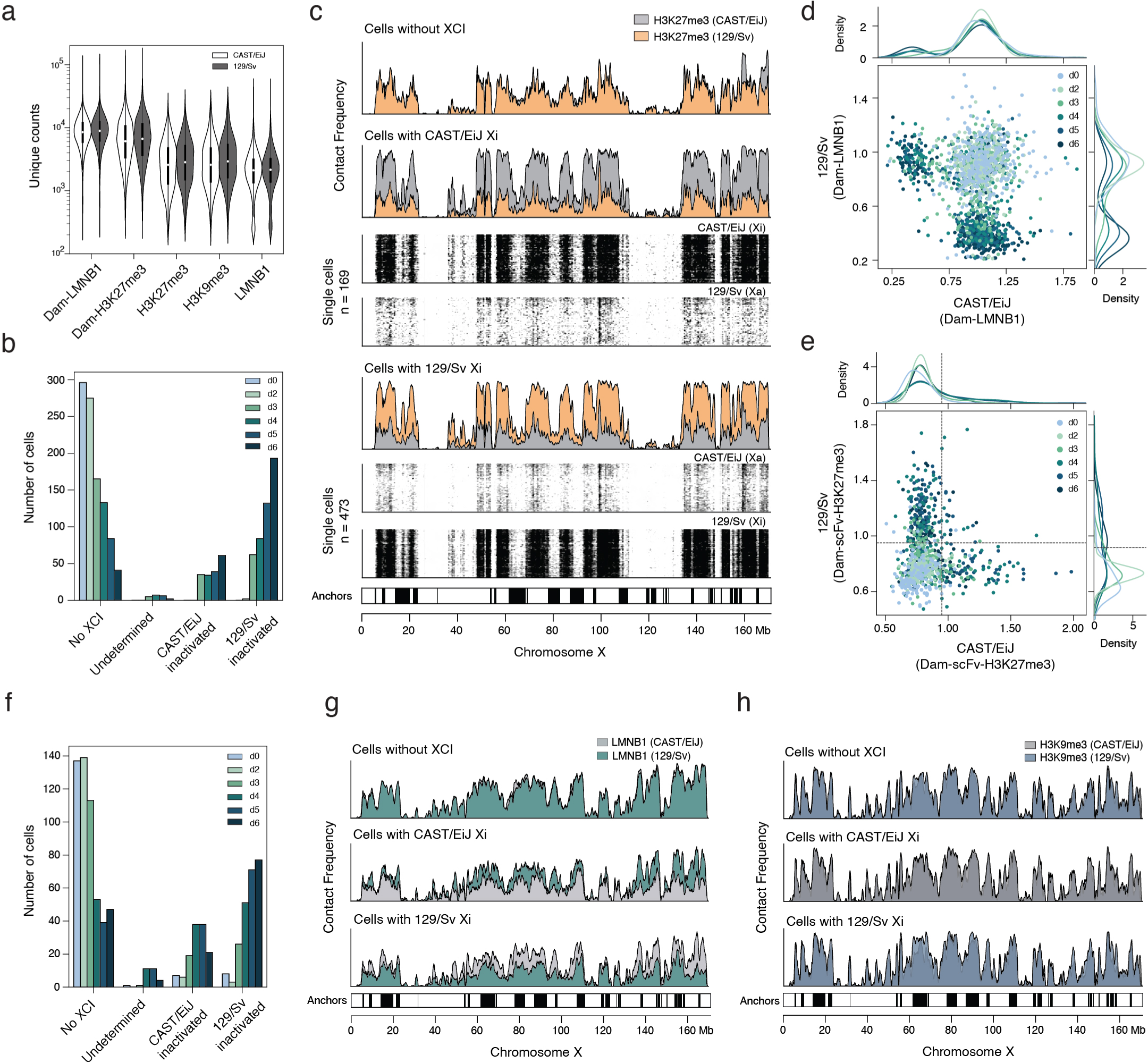
(**a**) Violin plots depicting the number of unique counts per cell and allele for the various readouts obtained with Dam&ChIC in the Vitamin C differentiation trajectory. (**b**) Number of cells in each Vitamin C differentiation day for all categories of cells based on allelic H3K27me3 levels in (4c). (**c**) Contact Frequency values of H3K27me3 on either allele for the cell categories defined in (4c), along the entire X chromosome. For the categories that undergo XCI, single-cell heatmaps of the active (Xa) and inactive (Xi) X chromosome are shown below, with the respective cell numbers. Bottom bar shows calls of the “anchors”. (d) Scatterplot of allelic Dam-LMNB1 levels per single cell in the Dam-LMNB1/H3K27me3 dataset, colored by differentiation day. (**e**) Scatterplot of allelic Dam-scFv-H3K27me3 levels per single cell in the Dam-scFv-H3K27me3/LMNB1 dataset, colored by differentiation day. The dashed lines indicate the manually set threshold used to classify cells as no XCI, CAST/EiJ inactive or 129/Sv inactive. (**f**) Number of cells in each Vitamin C differentiation day for all categories of cells based on Dam-scFc-H3K27me3 as in (S4e). (**g**) Contact Frequency values of LMNB1 on either allele for the cell categories defined in (S4e), along the entire X chromosome. The LMNB1 readout derives from the Dam-scFv-H3K27me3/LMNB1 dataset. (**h**) Contact Frequency values of H3K9me3 on either allele for the cell categories defined in (S4d), along the entire X chromosome. The H3K9me3 readout derives from the Dam-LMNB1/H3K9me3 dataset.

**Figure S5.**
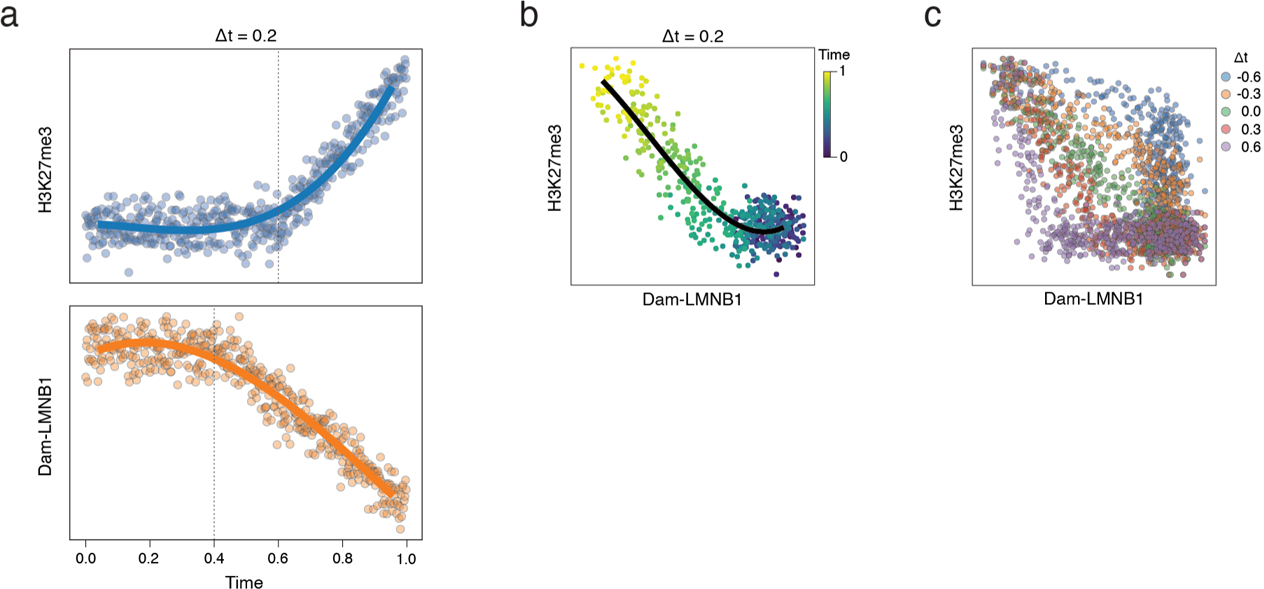
(**a**) Simulated single-cell H3K27me3 (top) and Dam-LMNB1 (bottom) values, where the onset of both dynamics is controlled to enable comparison of their relative timing. Values are generated by randomly sampling from a log-normal distribution and the onset of dynamics is simulated by randomly sampling from a Poisson distribution. Simulated time difference (Δt) is 0.2. (**b**) Scatterplot of simulated Dam-LMNB1 (x-axis) and H3K27me3 (y-axis) values from (a). Cells are colored according to simulated time. (**c**) Same as (b), but with cells following and colored according to progressive Δt values.

**Figure S6.**
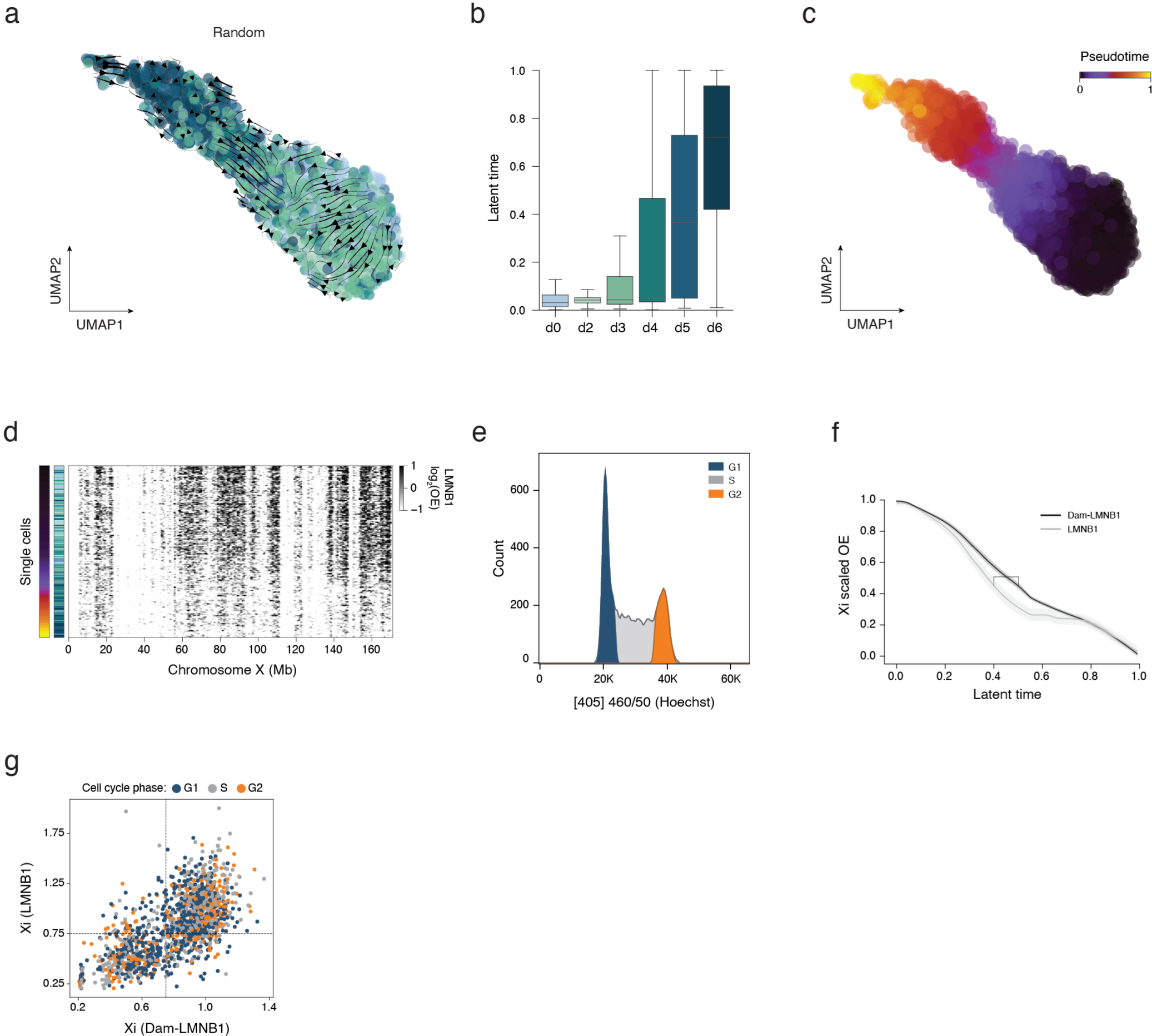
(**a**) Same UMAP as in (6a) based on Dam-LMNB1 levels of the inactive X in 100Kb bins, with streamlines representing chromatin velocities of randomly coupled Dam-LMNB1 and LMNB1 cells. (**b**) Distribution of latent time defined in (6b) across progressive differentiation days. (**c**) Same UMAP as in (S6a), cells colored according to pseudotime inferred from underlying UMAP using scVelo. (**d**) LMNB1 single-cell heatmap of the entire inactivating X-chromosome, ordered along the inferred latent time defined in (6b). (**e**) Hoechst profile from the Dam-LMNB1/LMNB1 Vitamin C experiment, showing the DNA content of cells and their respective cell cycle stage. (**f**) Average Dam-LMNB1 (black) and LMNB1 (grey) levels of the inactive X chromosome along latent time. Gap marks the moment of highest inactive X genome-lamina dynamics. (**g**) Same scatterplot as in (6g), with points colored according to cell-cycle stage defined by Hoechst.

